# Highly Multiplexed Spatial Transcriptomics in Bacteria

**DOI:** 10.1101/2024.06.27.601034

**Authors:** Ari Sarfatis, Yuanyou Wang, Nana Twumasi-Ankrah, Jeffrey R. Moffitt

## Abstract

Single-cell decisions made in complex environments underlie many bacterial phenomena. Image-based transcriptomics approaches offer an avenue to study such behaviors, yet these approaches have been hindered by the massive density of bacterial mRNA. To overcome this challenge, we combine 1000-fold volumetric expansion with multiplexed error robust fluorescence *in situ* hybridization (MERFISH) to create bacterial-MERFISH. This method enables high-throughput, spatially resolved profiling of thousands of operons within individual bacteria. Using bacterial-MERFISH, we dissect the response of *E. coli* to carbon starvation, systematically map subcellular RNA organization, and chart the adaptation of a gut commensal *B. thetaiotaomicron* to micron-scale niches in the mammalian colon. We envision bacterial-MERFISH will be broadly applicable to the study of bacterial single-cell heterogeneity in diverse, spatially structured, and native environments.

## Introduction

Population-level bacterial dynamics often emerge from the heterogeneous behaviors of single cells. Notable examples include entry into and exit from antibiotic persistent states (*1*), bet hedging (*2*), virulence factor expression (*3*), and cellular specialization within biofilms (*4*). Recent advances in bacterial single-cell RNA-sequencing (scRNA-seq) offer an exciting avenue to study such phenomena by providing transcriptome-wide expression profiles for thousands of cells (*5*– *14*). Indeed, such methods have provided insights into antibiotic response (*11*, *13*, *14*), prophage activation (*9*, *11*, *14*), toxin expression (*9*, *10*), sporulation (*10*), competence (*9*, *10*), mobile genetic elements (*11*, *14*), cell-cycle-dependent gene regulation (*15*), and functional heterogeneity within the rumen microbiome (*16*).

Missing from these studies is the natural spatial context in which many behaviors occur. Yet, spatial organization, across a range of length scales, is an essential modulator of bacterial dynamics. On the tens-of-micron-scale, spatial gradients in small molecule concentrations tune bacterial responses, define niches for commensal growth, or shape interactions in multi-species communities (*17*, *18*). On the micron-scale, direct cell-to-cell contact controls effector protein delivery which mediates predation (*19*), self-versus non-self recognition (*20*), contact-dependent inhibition (*21*), and virulence (*22*). Finally, even sub-micron length scales are relevant, as growing evidence indicates that the bacterial transcriptome is internally organized with functional consequences (*23*). Unfortunately, such spatial information is lost during cell dissociation and RNA extraction in scRNA-seq; thus, current methods are not well suited for the study of such processes.

By contrast, image-based approaches to single-cell transcriptomics provide this spatial context by directly imaging and identifying RNAs within fixed cells in their native spatial environment (*24*– *26*). Moreover, by leveraging combinatorial optical barcodes to distinguish RNAs, these measurements can be massively multiplexed, producing spatially resolved, transcriptome-scale expression profiles that span intracellular to tissue-scale lengths (*24–26*). In eukaryotic systems such methods have mapped the intracellular RNA organization, explored regulatory networks, and defined, discovered, and charted cell types and states across a range of tissues (*24–26*). Unfortunately, current methods are not compatible with bacteria, as the massive density of bacterial RNA challenges combinatorial barcode detection. Non-combinatorial barcoding approaches can bypass this challenge, as recently illustrated with *Pseudomonas aeruginosa* (*27*), yet are not suitable for transcriptome-wide profiling.

Here we overcome this RNA density challenge and introduce a transcriptome-scale, image-based approach for bacterial, single-cell transcriptomics. This approach combines a bacterially optimized expansion microscopy toolbox with multiplexed error-robust fluorescent *in situ* hybridization (MERFISH) (*28*) and allows single-cell profiling of up to 80% of the transcriptome. We demonstrate that this technique—bacterial-MERFISH—accurately profiles 97, 1,057, or 1,930 operons with large detection efficiency, accuracy, and throughput in log-phase *Escherichia coli* (*E. coli*) cells. To highlight the discovery potential of this technique, we first profile *E. coli* response to a carbon-source switch, revealing a heterogeneous sequential nutrient exploration program. Next, we chart the intracellular organization of the *E. coli* transcriptome, uncovering a previously unappreciated diversity in spatial patterning and a cooperative role for genome and proteome organization in shaping transcriptome organization. Finally, we map the adaptation of a human gut commensal—*Bacteroides thetaiotaomicron* (*B. theta*)—to the mouse colon, revealing micron-scale fine-tuning of gene expression based on local polysaccharide availability. More broadly, these measurements illustrate the potential for bacterial-MERFISH to reveal single-cell heterogeneity in a wide range of biological and spatial contexts.

## Results

### Bacterial-MERFISH accurately profiles thousands of operons in E. coli

MERFISH enables the identification of thousands of different mRNA molecules by using combinatorial, error-robust, fluorescent optical barcodes built from repetitive rounds of single-molecule FISH (smFISH) (*28*). However, to decipher barcodes, the fluorescent signal from different molecules must be optically resolvable. For conventional high-resolution optical microscopy, only a few molecules per µm^3^ can be distinguished (*29*). For eukaryotic systems, large transcriptome fractions can be targeted while satisfying this limit (*28*). By contrast, a log-phase *E. coli* cell contains ∼8,000 mRNA molecules in a cell volume of ∼3 µm^3^ (*30*), producing a total mRNA density nearly three orders of magnitude greater than that resolvable with diffraction-limited imaging (Fig. 1A). Thus, mRNA density restricts the imaging of more than a small number of bacterial mRNAs (*31*) and is a substantial challenge to transcriptome-scale imaging.

**Fig. 1.**
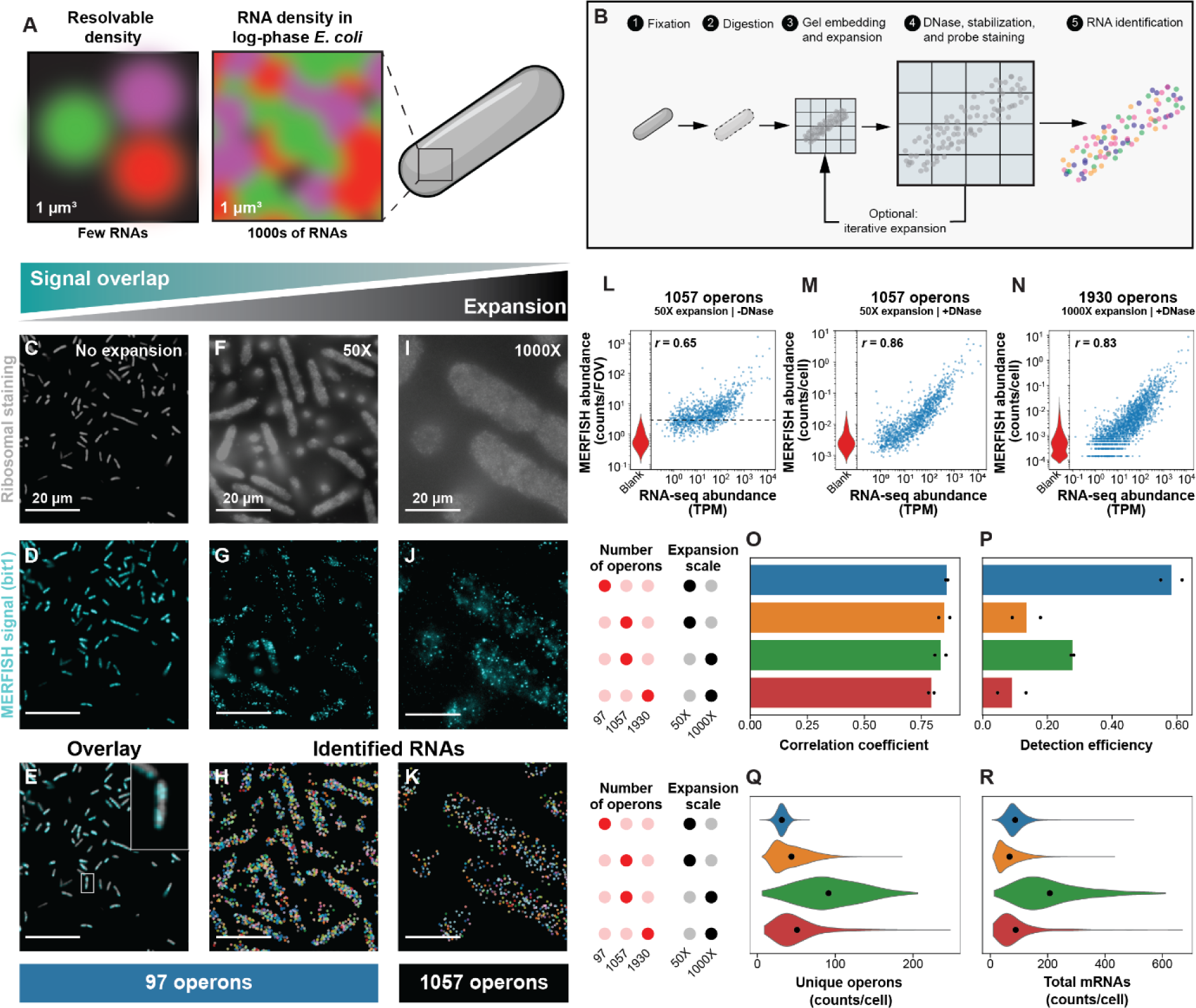
Image-based expression profiling of thousands of operons in *E. coli* with bacterial-MERFISH. (**A**) The RNA density resolvable with fluorescence microscopy and the estimated *E. coli* mRNA density. (**B**) Protocol for bacterial-MERFISH. (**C-E**) Images of fixed, log-phase *E. coli* grown in Luria Broth (LB) stained for the 16S ribosomal RNA (rRNA) (C), the first bit of a 97-operon MERFISH measurement (D), or overlay of the two (E). Scale bars: 20 µm. (**F,G**) As in (C,D) but for 50X-expanded *E. coli*. (**H**) RNA identity (color) determined by MERFISH for cells in (F,G). (**I-K**) As in (F-H) but for 1000X-expanded *E. coli* stained with a 1,057-operon library. (**L**) Average mRNA copy number per field of view (FOV) for a 50X-expanded, 1,057-operon MERFISH measurement without DNase treatment versus bulk RNA-sequencing (RNA-seq) abundance from a matched culture. Dashed line: false-positive rate estimated from the average MERFISH abundance for the 100 mRNAs with the lowest RNA-seq abundance. TPM: transcripts per million reads. Left: distribution of MERFISH false-positive-controls (‘blank’). (**M,N**) As in (L) but for 50X-expanded, 1,057-operon MERFISH (M) or 1000X-expanded, 1,930-operon MERFISH (N) with DNase treatment. *r:* Pearson correlation coefficient between logarithmic expression. (**O,P**) Correlation coefficient with bulk sequencing (O) or detection efficiency (P) for different MERFISH measurements. Bars represent the average between two replicates (markers). (**Q,R**) Distribution of unique operons (Q) or total mRNAs (R). Markers represent the average.

Density reduction is a natural avenue to address this challenge. Indeed, image-based approaches to single-cell transcriptomics have achieved modest degrees of RNA density reduction by spreading the barcode signal over more imaging rounds (*32*, *33*) or by leveraging expansion microscopy (*34*) to physically swell the sample (*32*, *35*). While the approximately 10-fold reduction in RNA density achieved by these approaches was sufficient to extend image-based transcriptomes to whole-transcriptome-scale in eukaryotes (*32*, *33*), such density reduction is still nearly two orders of magnitude insufficient for similar profiling in bacteria.

To overcome bacterial mRNA density, we leveraged recent advances in expansion microscopy (*36*, *37*) to develop a bacterial-FISH-optimized expansion toolbox capable of up to 1000-fold volumetric expansion (Fig. 1B), complementing recent bacterial expansion methods developed for non-RNA targets and with modest degrees of expansion (*38–43*). Briefly, we grew *E. coli* to mid-log phase, fixed them with paraformaldehyde (PFA), digested the cell wall, expanded them in a Ten-fold Robust Expansion (TREx) gel (*36*), and then re-embedded the sample in a non-expanding, stabilizing gel (Fig. 1B; Materials and Methods). Using custom expansion-optimized staining protocols, we labeled samples with a 16S ribosomal RNA (rRNA) probe and a MERFISH probe set targeting 97 *E. coli* operons (Materials and Methods). We targeted operons rather than individual genes as the signals from different barcodes from genes on the same polycistronic mRNA would not be resolvable.

Expansion increased the width of *E. coli* 3.9±0.9-fold (standard deviation [STD], *n*=4; fig. S1A) consistent with a ∼50-fold volumetric expansion (fig. S1B). Thus, we term this approach the 50X expansion protocol. In expanded cells stained with the 97-operon library, individual fluorescent puncta were visible, and these molecules were identified with MERFISH (Fig. 1, C to H), indicating sufficient expansion to resolve the mRNA density of this targeted library. Interestingly, we noted ample number of molecules even in the absence of specific RNA-gel anchoring chemistries in PFA-fixed but not methanol-fixed expanded samples, suggesting a PFA-dependent, RNA-anchoring mechanism (fig. S1, C to I).

To further explore the multiplexing possible with the 50X protocol, we stained 50X-expanded *E. coli* with a MERFISH probe set targeting 1,057 operons, roughly 40% of the *E. coli* transcriptome (Materials and Methods). While the mRNA density was higher than that observed for the 97-operon measurement, individual puncta were still resolved and many molecules were identifiable with MERFISH (fig. S1, J to L). Nonetheless, we noticed an increased frequency of overlapping RNA signals (fig. S1K). To address this overlap, we developed an iterative expansion protocol that combined TREx (*36*) with a previous iterative strategy (*37*). Briefly, cells expanded and stabilized with the 50X protocol were expanded in a second TREx gel and then embedded in a second stabilizing gel (Fig. 1B; Materials and Methods). This protocol expanded the width of cells 11.1±1.7-fold (STD, *n=*4; fig. S1A), consistent with a ∼1,300-fold volumetric expansion (fig. S1B). Thus, we term this protocol the 1000X protocol. When cells were expanded with this protocol, the signal from individual RNAs and their identity were clearly distinguished when 1,057 operons were stained (Fig. 1, I to K). Notably, the 1000X protocol starts with 50X-expanded samples (Fig. 1B), facilitating exploration of the necessary expansion for a given sample.

Inspired by the ability to expand *E. coli* volumetrically by three orders of magnitude, we designed a MERFISH library that covers 80% of the transcriptome, corresponding to 1,930 operons (Materials and Methods). 1000X-expanded samples stained with this probe set showed clear single-molecule signals with limited overlap, and these RNAs could be identified with MERFISH (fig. S1, M to O), suggesting that 1000X expansion is sufficient to allow MERFISH profiling of a substantial fraction of the *E. coli* transcriptome.

Notably, during the early development of bacterial-MERFISH, we observed a low correlation between the abundance determined via bacterial-MERFISH and that of bulk RNA-sequencing for lowly expressed operons, suggesting a higher false-positive rate than that measured by our internal false-positive controls (Fig. 1L; Materials and Methods). This discrepancy suggested a bacterial-expansion-dependent source of false positives, which we reasoned might be due to probe binding to genomic DNA melted during expansion. Supporting this hypothesis, DNase treatment dramatically reduced this apparent false-positive rate, bringing it into agreement with that measured with internal controls (Fig. 1M).

To benchmark the performance of bacterial-MERFISH, we performed two replicate measurements of 97, 1,057, or 1,930 operons in combination with 50X or 1000X expansion in log-phase *E. coli*, segmented cells from these images, and partitioned RNAs into those cells (Fig. 1, F to K, and fig. S1, J to O). We observed strong correlation between these measurements and bulk RNA-sequencing across all conditions (Fig. 1, M to O, and fig. S2, A and B), indicating that bacterial-MERFISH can accurately profile RNA expression across four orders of magnitude in abundance. Further supporting its accuracy, measurements with bacterial-MERFISH correlated strongly between biological replicates, expansion protocols, and multiplexing levels (fig. S2, C to E).

These measurements also revealed that the detection efficiency—the fraction of targeted molecules actually detected—of bacterial-MERFISH is comparable to or greater than the capture efficiencies reported for scRNA-seq (*5–14*) (Fig. 1P and fig. S2F; Materials and Methods). Values ranged from 7% for near-whole transcriptome measurements (1,930 operons) to as high as 50% for more targeted measurements (97 operons), with additional expansion providing an increase from 10% to 25% for 1,057 operons measurements (Fig. 1P). As expected, the number of mRNA counts and unique operons observed per cell varied based on the multiplexing and detection efficiency (Fig. 1, Q and R). Finally, bacterial-MERFISH can image large numbers of cells, comparable or greater than those characterized previously with scRNA-seq (*5–16*), despite the decreased imaging throughput due to expansion (fig. S2G). Importantly, with the ability to vary multiplexing and expansion, it is possible to balance different aspects of performance, e.g., the number of imaged cells versus the degree of expansion, to best suit the question. Collectively, these measurements indicate that bacterial-MERFISH is a high-performance, versatile image-based approach to bacterial single-cell transcriptomics, with low false-positive rates and large dynamic range, detection efficiencies, and throughput.

### Bacterial-MERFISH reveals an asynchronous hierarchical response during a shift in carbon source

One promise of single-cell bacterial transcriptomics is the ability to identify heterogeneous bacterial behaviors and to computationally resynchronize the desynchronized response of individual cells to environmental stimuli, unmasking dynamics obscured by population averages. One illustrative dynamic response is the diauxic shift caused by a switch in carbon source. Bacteria often consume different carbon sources in a preferential order regulated, in part, by carbon-catabolite repression (CCR) (*44*). A bacterial population grown in a mixture of two carbon sources will often consume the preferred source, pause growth in a diauxic shift, and then resume growth on the less preferred substrate (*45*). The diauxic shift is classically interpreted as the time required to express utilization machinery for the second sugar; however, recent single-cell studies suggest that this response is instead shaped by differential dynamics of sub-populations (*46*). Nonetheless, the diversity and transcriptional profiles of such sub-populations remain poorly defined.

Thus, we revisited the classic diauxic shift with bacterial-MERFISH. We grew *E. coli* in a minimal defined medium with a mixture of glucose and xylose (Materials and Methods). As expected, the culture grew logarithmically until glucose was exhausted, paused growth in a diauxic shift, and then resumed logarithmic growth on xylose before entering stationary phase once both sugars were exhausted (Fig. 2A). We harvested *E. coli* cells throughout this process and profiled the expression of 1,057 operons (Fig. 2, A to C) in 296,666 cells with 50X expansion—a multiplexing and expansion level set to balance transcriptome-wide profiling with the number of profiled cells. For experimental efficiency, multiple time points were combined and profiled in a single MERFISH measurement by labeling cells with barcoded 16S rRNA probes (*27*) optimized for expansion protocols (Fig. 2B; Materials and Methods). Putative cells were segmented, RNA molecules assigned to these cells, and conditions demultiplexed with the 16S rRNA barcode (Fig. 2C; Materials and Methods).

**Fig. 2.**
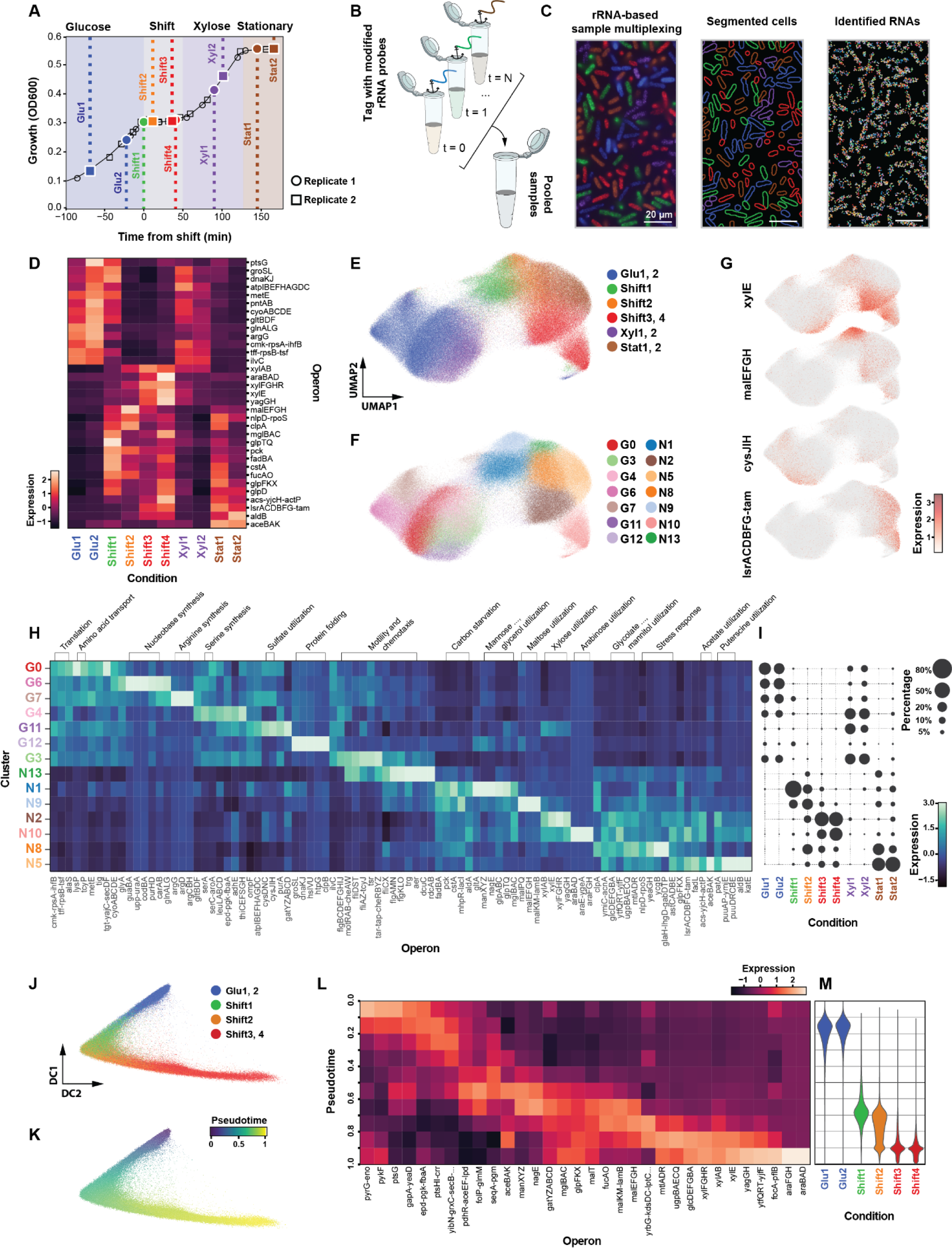
Bacterial-MERFISH reveals a stochastic hierarchical response to carbon starvation. (A) Optical density (OD) at 600 nm of *E. coli* growing in minimal defined medium containing a mixture of glucose and xylose. Markers: biological replicates, colored if sampled for MERFISH. 16S rRNA sample multiplexing strategy. (**C**) rRNA image colored by identified sampling condition (left), segmented cell boundaries similarly colored (middle), and identified RNAs (color; right) for a 50X-expanded, 1,057-operon MERFISH measurement. rRNA colors are defined as in (A). Scale bars: 20 µm. (**D**) Average gene expression measured in z-score for sampling conditions as listed in (A). (**E-G**) UMAP of cells colored by condition (E), Leiden cluster (F), or the natural log of normalized gene expression (G). Clusters are numbered in order of abundance and labeled based on growth (*‘G’*) or non-growth (*‘N’*). (**H,I**) Marker gene expression measured in z-score (H) and relative fraction in each condition (I) for clusters. (**J,K**) Scatter plot of the first two diffusion components for cells collected from glucose log-phase or diauxic shift colored by condition (J) or pseudotime (K). (**L**) Average carbon-utilization operon expression measured in z-score at different pseudotimes. (**M**) Distribution of the pseudotime values by condition.

Supporting these measurements, we found that the average RNA expression determined via MERFISH was strongly correlated between biological replicates and with bulk RNA-sequencing at different growth stages (fig. S3, A to D). Moreover, cell populations displayed expected expression patterns (Fig. 2D) with cells harvested during growth expressing operons associated with amino acid synthesis (e.g., *argG* and *ilvC*), translation (e.g., *tff-rpsB-tsf* and *cmk-rpsA-ihfB*), and aerobic respiration (e.g., *atpIBEFHAGDC*), and cells harvested during the diauxic shift and stationary phase expressing operons associated with stress response (e.g., *nlpD-rpoS*) and gluconeogenesis (e.g., *glpFKX* and *glpD*).

To explore the heterogeneity in cellular response to this carbon shift, we integrated the measurements from two biological replicates, visualized single-cell heterogeneity with Uniform Manifold Approximation and Projection (UMAP), and performed Leiden clustering to distinguish sub-populations (Fig. 2, E to I; Materials and Methods). Supporting our analysis, cells harvested at similar growth phases largely co-integrated (fig. S3E), yet, despite modest transcriptional differences, cells collected from log-phase growth in glucose or xylose were largely resolved (Fig. 2E and fig. S3F). This analysis revealed a rich diversity in behavior at the single-cell level. Cells were organized into two major groups, corresponding to cells taken from conditions of growth or non-growth (Fig. 2E), and were collectively sub-divided into 14 different clusters (Fig. 2F). These clusters had unique gene expression profiles (Fig. 2, G and H) and abundances across different conditions (Fig. 2I). Supporting cluster validity, individual clusters were observed across both replicates (Fig. 2I) and were each marked by multiple operons associated with related biological processes (Fig. 2H). Moreover, single-molecule FISH (smFISH) in unexpanded *E. coli* confirmed the co-expression of cluster markers (fig. S4).

During log-phase growth, we identified a diversity of sub-populations (Fig. 2, E to I), including clusters associated with translation and amino acid transport (G0 marked by *tff-rpsB-tsf* and *lysP*), nucleobase synthesis (G6 marked by *carAB*, *codBA*, and *purHD*), arginine synthesis (G7 marked by *argG*, *argD*, and *argCBH*), serine biogenesis (G4 marked by *serA* and *serC-aroA*), sulfate utilization (G11 marked by *cysDNC* and *cysJIH*), protein folding (G12 marked by *dnaKJ*, *groSL*, *hslVU*, *clpB*, and *htpG*), and motility and chemotaxis (G3 marked by *motRAB-cheAW*, *fliDST*, and *tar-tap-cheRBYZ*) (Fig. 2H). Intriguingly, many of these biochemical processes are required for growth in minimal media, yet our analysis revealed that only subsets of cells expressed high levels of these essential operons, suggesting a model in which the homeostatic, population-level expression of these pathways is produced not by uniform expression across all cells but rather by transient, coherent bursts of expression that time-average to required levels. This observation is consistent with transient bursts in promoter activity revealed with live-cell imaging (*47*, *48*) and the presence of similar clusters in recent scRNA-seq measurements (*10*). Our measurements now suggest that such bursts are widespread and occur not at the individual operon level but rather upstream in common regulatory factors.

We also observed a similar degree of heterogeneity during the diauxic shift (Fig. 2, E to I). Interestingly, only a subset of clusters expressed operons associated with xylose utilization (N2 and N10 both marked by *xylAB*, *xylE*, *xylFGHR*, and *yagGH*) while most clusters in the diauxic shift expressed utilization operons associated with carbon sources not included in our medium (Fig. 2, E to I). These sources comprised mannose and glycerol (N1 marked by *manXYZ*, *glpABC*, and *glpTQ*); maltose (N9 marked by *malEFGH*, *malKM-lamB*, and *malPQ*); arabinose (N10 marked by *araE-ygeA*, *araFGH*, and *araBAD* in addition to xylose-associated operons); mannitol and glycolate (N8 marked by *mtlADR* and *glcDEFGBA*); and acetate and puterscine (N5 marked by *acs-yjcH-actP*, *puuAP-ymjE*, and *puuDRCBE*) (Fig. 2, H and I). N13, like G3, was marked by chemotaxis and motility operons. However, these clusters were distinguished by operons associated with chemotaxis towards peptides (G3; e.g., *tar-tap-cheRBYZ* and *tsr*) or sugars (N13; e.g., *trg* and *aer*), reflecting the differential nutrient needs between the conditions associated with these clusters (Fig. 2H).

We also noted an apparent temporal order in which specific carbon sources were explored. To investigate this ordering, we computationally resynchronized cells using a pseudotime analysis from glucose-log-phase growth through the diauxic shift (Materials and Methods). Not only did this analysis reproduce the order of sampling during the diauxic shift (Fig. 2, J and K), supporting the pseudotime ordering, it also revealed a temporal cascade of carbon-utilization operon expression (Fig. 2L). This cascade started with operons associated with glucose (e.g., *ptsG*) and then proceeded through operons associated with glycogen (e.g., *segA-pgm*); the glyoxylate cycle— related to acetate utilization—(e.g., *pdhR-aceEF-lpd* and *aceBAK*); mannose (e.g., *manXYZ*); galactitol and galactose (e.g., *gatYZABCD* and *mglBAC*); glycerol (e.g., *glpFKX*); maltose and fucose (e.g., *malT*, *malEFGH*, *malKM-lamB*, and *fucAO*); xylose (e.g., *xylFGHR* and *xylAB*); and then, finally, arabinose (e.g., *araFGH* and *araBAD*). The transient co-expression of xylose- and arabinose-utilization operons seen at late pseudotime values (Fig. 2L) may represent an overshoot in carbon-source progression due to the lag between transcription of xylose-utilization operons and the transition to the functional utilization of xylose. Importantly, the range of pseudotime values observed for each shift condition overlapped with those of other conditions (Fig. 2M), supporting a stochastic response model in which cells are desynchronized in their progression along this carbon-source hierarchy. Notably, this model does not assume that all cells explore all carbon sources.

Together these results reveal that the homeostatic levels of required pathways can be maintained by transient, coherent bursts of expression within entire regulons, and that, when faced with carbon starvation, *E. coli* adopts a responsive diversification strategy (*49*) in which the lack of glucose triggers a stochastic hierarchical progression along sugar utilization operons. Importantly, with the ability to characterize sub-populations and computationally resynchronize cells undergoing dynamic responses, bacterial-MERFISH could prove useful in dissecting the diverse regulatory mechanisms that encode such cellular heterogeneity.

### Bacterial-MERFISH reveals a diversity of intracellular localization patterns for E. coli mRNAs

Another advantage of image-based, single-cell transcriptomic methods is the ability to measure the intracellular transcriptome organization. Specific bacterial mRNAs are localized to the cytoplasm (*50–52*), membrane (*50–52*), poles (*50*, *52–55*), chromosomal loci of origin (*56*), or the surface of intracellular organelles (*57*) as determined via low throughput mRNA imaging (*50–57*) or biochemical fractionation (*50*, *52*). This organization has proposed functional roles in protein sorting, complex assembly, and mRNA turnover (*23*). However, as RNA localization has yet to be imaged at the transcriptome-scale, the extent and diversity of spatial organization remain unclear.

To explore the spatial organization of the *E. coli* transcriptome, we leveraged our 50X-expanded, 1,057-operon measurements of *E. coli* grown in LB. To define intracellular RNA localization, we mapped each mRNA molecule to its axial and radial position within each cell and normalized these coordinates to the cell length and width (Fig. 3A; Materials and Methods). We then computed the average axial and radial distributions for each mRNA across all measured cells (Fig. 3, B to D; Materials and Methods). Consistent with previous reports in *E. coli* (*23*), we identified mRNAs enriched in the cytosol (e.g., *dnaKJ*), at the membrane (e.g., *ptsG*), and toward the poles (e.g., *dnaB*) (Fig. 3, C and D). However, we also noticed a striking diversity of variations on these major patterns, with mRNAs enriched at different locations along the membrane, at multiple cytoplasmic locations, or in a single central focus (Fig. 3E). These patterns were reproduced between replicate measurements (fig. S5, A to C), with 1000X-expanded MERFISH measurements (fig. S5D), and with unexpanded smFISH (Fig. 3F).

**Fig. 3.**
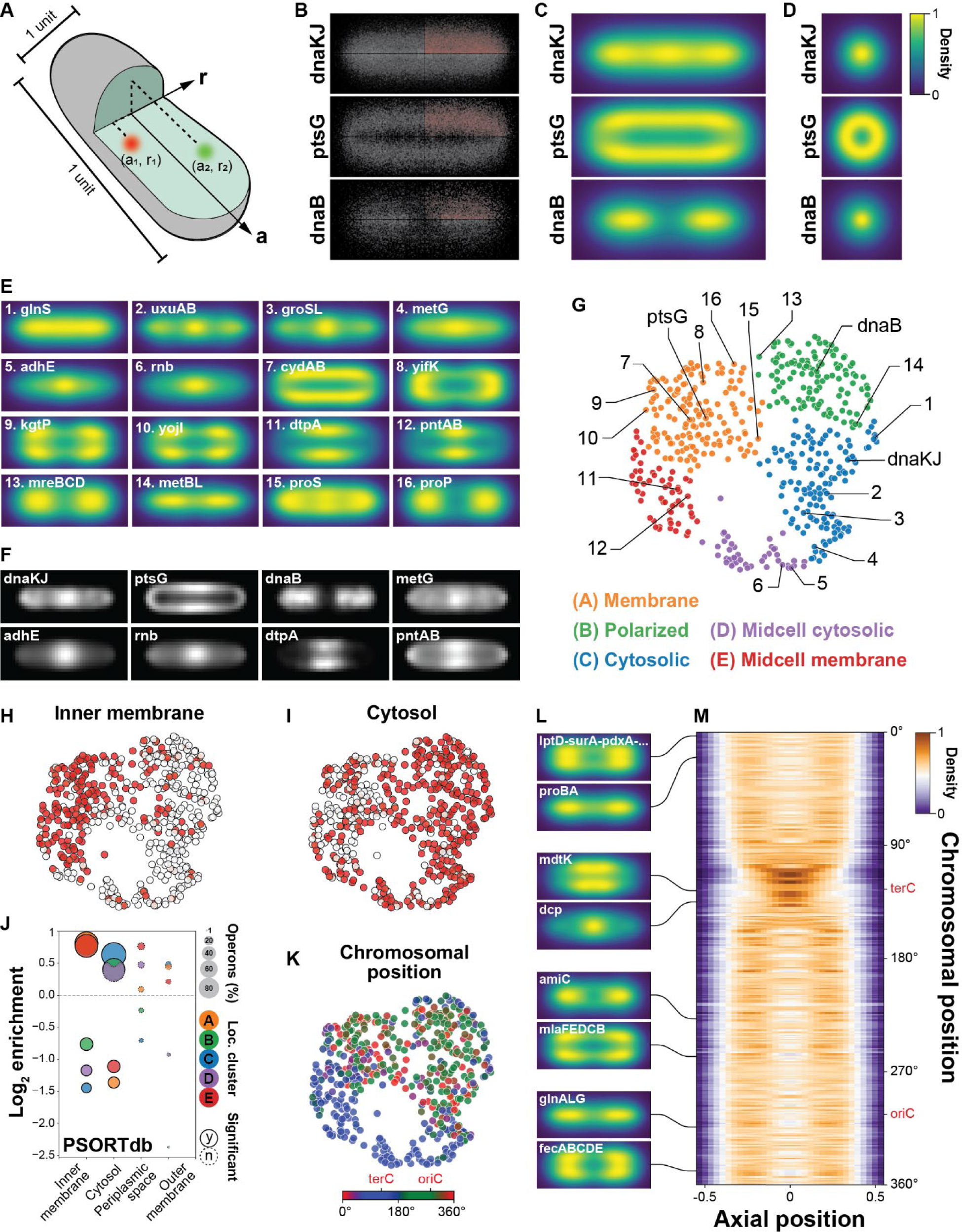
The *E. coli* transcriptome is organized in diverse patterns shaped by proteome and genome organization. (**A**) Mapping mRNAs to a normalized cellular coordinate system. (**B**) Location of *dnaKJ* (top), *ptsG* (middle), or *dnaB* (bottom) mRNAs measured with 50X-expanded, 1,057-operon MERFISH in LB log-phase *E. coli*. mRNAs were mapped to a positive axial midcell distance and radial coordinate (red upper quadrant) and mirrored (gray quadrants) for visualization and analysis. (**C,D**) Smoothed, normalized RNA density for the localizations plotted in (B) along the radial and axial directions (C) or radial directions (D). (**E**) As in (C) for an illustrative mRNA set. (**F**) Averaged, normalized RNA signal for unexpanded smFISH of individual operons for a single z-plane. (**G**) UMAP of mRNA spatial distributions colored by cluster and marked by name or number for examples in (C) or (E), respectively. (**H,I**) UMAP as in (G) colored by predicted location, inner membrane (H) or cytoplasm (I), of the encoded proteins for each mRNA. (**J**) Enrichment of protein location label within each cluster. Color: cluster. Size: fraction of operons with label. Boundary: significance of enrichment (‘y’ is an FDR-corrected p-value less than 0.05). (**K**) UMAP as in (G) colored by chromosomal position from which each operon is transcribed. The origin (*oriC*) and terminus (*terC*) of replication are listed. (**L**) As in (C) for example mRNAs. (**M**) Normalized, averaged, axial mRNA density arrayed by the chromosomal location from which they were transcribed.

To further categorize this spatial diversity, we leveraged a measure of pattern similarity to visualize and cluster spatial patterns (Materials and Methods). This analysis produced five major clusters of mRNA distributions (Fig. 3G) which showed some degree of continuous spatial variation within the clusters. The Cytoplasmic cluster comprised patterns including uniform filling of the cytoplasm (e.g., *dnaKJ* and *glnS*), multiple foci throughout the cell (e.g., *uxuAB* and *groSL*), or increased central density (e.g., *metG*). mRNAs with a strong central focus (e.g., *adhE* and *rnb*) defined the Midcell-cytosolic cluster. The Membrane cluster was defined by diffuse membrane-enriched patterns (e.g., *ptsG* and *cydAB*) or membrane enrichment at or adjacent to the poles (e.g., *yifK*, *kgtP* or *yojI*) while the Midcell-membrane cluster was defined by strong membrane enrichment only in the middle of the cell (e.g., *dtpA* and *pntAB*). Finally, the Polarized cluster was defined by diffuse (e.g., *mreBCD*) or sharp cytoplasmic foci (e.g., *metBL*) near the poles. Importantly, some mRNAs at cluster boundaries shared features similar to nearby clusters (e.g., *proS* and *proP*), underscoring a continuous variation in spatial patterns.

To investigate possible patterning mechanisms, we explored the correlation between clusters and mRNA features. Covariation with spatial distribution was modest for transcript length, GC content, abundance, and half-life (fig. S6, A to H) but strong for encoded protein location (Fig. 3, H to J, and fig. S6, I to N). Specifically, operons containing mRNAs that encode at least one inner-membrane protein were enriched in membrane-associated RNA localization clusters, whereas mRNAs that encode cytoplasmic proteins were enriched in clusters found within the cytosol (Fig. 3J). These observations are consistent with previous measurements for individual mRNAs (*50*) as well as transcriptome-wide mRNA groups (*51*); with the co-translational insertion mechanism of inner-membrane proteins, which would concentrate mRNAs at the membrane during translation (*58*); and with the report of membrane-associated sequence features enriched in inner-membrane-protein-coding mRNAs (*59*, *60*).

In parallel, the genomic locus from which the mRNA was transcribed also correlated strongly with spatial patterning. mRNAs within the Midcell-cytosolic or Midcell-membrane clusters were preferentially encoded from genomic loci near the terminus of replication (*terC*) while the genes for mRNAs in other clusters were depleted in this chromosomal region (Fig. 3K and fig. S6O). During conditions of fast growth, the *E. coli* chromosome is organized with *terC* in the cell center and *oriC* and the left and right chromosomal arms replicating near the poles. This structure is maintained for the majority of the cell cycle, with the polar *terC* macrodomain formed at the pole of a newly divided cell moving rapidly to the cell center (*61*). Indeed, the axial distribution of mRNAs averaged across all (Fig. 3, L and M) or portions (fig. S6P) of the cell cycle were consistent with this organization.

Our measurements now unify two previous models suggested for bacterial transcriptome organization—that either genomic (*56*) or proteomic (*50–52*) features dictate organization. Specifically, we show that both features play a role in the global organization of mRNAs, with protein location shaping cytoplasmic versus membrane enrichment and genomic feature shaping axial enrichment both within the cytoplasm or on the membrane. Nonetheless, we identified multiple mRNAs with spatial patterns that deviate from these global rules, including membrane enriched mRNAs that do not encode known inner-membrane proteins (fig. S7, A and B), cytoplasmic-enriched mRNAs that encode an inner-membrane protein (fig. S7, C and D) and many mRNAs with spatial patterns inconsistent with those predicted for their genomic loci (fig. S7, E to G). These exceptions raise the possibility that there are mRNA-specific localization mechanisms that remain to be discovered and that these exceptions, in addition to global patterns, might have functional significance. Importantly, bacterial-MERFISH offers a direct approach to measuring such patterns, which should greatly enable mechanistic and functional studies of the intracellular organization of the bacterial transcriptome.

### Bacterial-MERFISH reveals adaptation of a gut commensal to distinct niches in the colon

Image-based approaches to single-cell transcriptomics also promise the ability to explore gene expression within complex, spatially structured environments. To explore this capability of bacterial-MERFISH, we leveraged germ-free mice monocolonized with the human gut commensal *B. theta* (Fig. 4A). As *Bacteroides* have the ability to harvest a remarkable diversity of polysaccharides—both from the rich dietary pool as well as those deposited onto the mucus layer by the host (*62*)—we reasoned that *B. theta* might modulate its polysaccharide utilization based on local polysaccharide availability, as suggested previously (*63*). To explore this possibility, we designed a MERFISH panel that covers the diverse importers (i.e., SusC proteins) within polysaccharide utilization loci (PUL), hybrid two-component systems (HTCS) often associated with the regulation of PUL expression, and a handful of genes associated with central metabolism and other bacterial functions, targeting 159 operons in total (Fig. 4B).

**Fig. 4.**
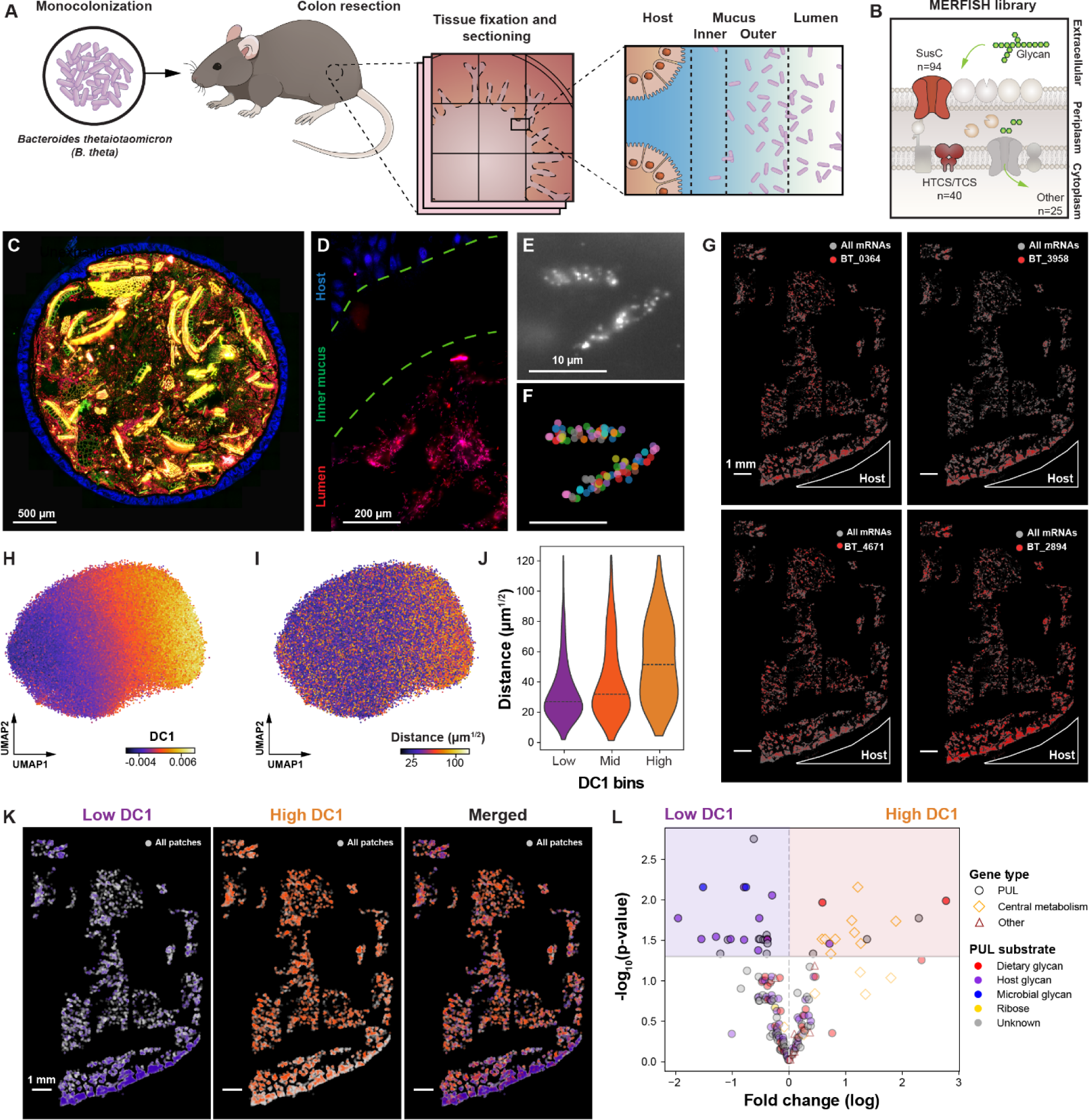
Adaptation of a gut commensal to micron-scale niches in the mammalian colon. (**A,B**) The colon from *B. theta* monocolonized mice was dissected, fixed, sliced, and expanded (A), and MERFISH targeting 159 operons—comprising polysaccharide utilization loci (PUL), hybrid two-component systems (HTCS), and other operons—was performed (B). (**C**) Unexpanded colon section colored by 16S rRNA (red), DAPI (blue), and dietary debris autofluorescence (green). Yellow indicates autofluorescence in multiple channels. Scale bar: 500 µm. (**D**) 50X-expanded colonic section colored by DAPI (blue) and the first bit of a MERFISH measurement (magenta). Scale bar: 200 µm. (**E,F**) The first bit of a MERFISH measurement (E) and RNA identity (F) for two *B. theta* cells imaged in (D). Scale bars: 10 µm. (**G**) Spatial distribution of all (gray) or individual RNAs (red). Host tissue is marked. Scale bars: 1 mm. (**H,I**) UMAP of patch gene expression colored by the first diffusion coefficient (DC1; H) or distance to host tissue (I). (**J**) Distribution of patch-host distance for patches with low, mid, and high DC1 values. Lines: median. (**K**) Spatial distribution of patches with low (blue), high (orange), or intermediate (gray) DC1 values for slice in (G). (**L**) Enrichment significance versus log-fold change for operon expression in high-versus low-DC1 patches. Marker: gene type. Marker fill color: PUL substrate, where known. The colored regions indicate an FDR-corrected p-values less than 0.05.

We then resected the colon from a monocolonized mouse, and the sample was methacarn-fixed, paraffin-embedded, and sectioned (Fig. 4A; Materials and Methods). In an unexpanded section, 16S rRNA staining revealed depletion of *B. theta* from the sterile inner mucus layer next to the host epithelial layer, enrichment near the outer mucus layer (*64*), and variable density throughout the colonic lumen (Fig. 4, A and C). We then expanded a section with a modified 50X-protocol and stained it for MERFISH. The 16S rRNA signal revealed that the distribution of *B. theta* was largely preserved despite a notable loss in dietary debris (Fig. 4, C and D; Materials and Methods). Critically, individual mRNAs could be identified by MERFISH in single *B. theta* cells (Fig. 4, E and F). Across this colonic cross section, we observed that the expression of many *B. theta* operons was spatially variable. Some mRNAs were expressed throughout the lumen (e.g., BT_0364 and BT_4671) while others were more prominently expressed near the mucus layer (e.g., BT_3958 and BT_2894) (Fig. 4G). Multi-color smFISH in unexpanded samples supported these expression patterns (fig. S8).

To quantify this spatial variation, we performed these measurements in six slices sampled from the colon of two mice. Each colon was fixed with one of two different methods to control for potential fixation-dependent artifacts (Materials and Methods), which were minor as evidenced by strong correlation between these measurements (fig. S9, A to C). As PUL expression can be low (fig. S8), we averaged gene expression over small spatial patches containing ∼5 *B. theta* cells and visualized transcriptional variation across patches using UMAP and diffusion maps (Fig. 4H and fig. S9, D to F). Individual genes expressed more prominently near the mucus layer or throughout the lumen defined different regions of this UMAP (fig. S9, D and E), and the diffusion analysis suggested that an important axis of variation, captured by the first diffusion component (DC1), is related to mucus proximity. Patches of low DC1 values were found enriched near the mucus whereas patches of high DC1 values were distributed more uniformly throughout the lumen (Fig. 4, H to K, and fig. S9F). While these observations reveal that mucus proximity is a major driver of *B. theta* expression variation, the presence of low-DC1 patches deeper into the lumen and many high-DC1 patches near the mucus, suggest that local spatial heterogeneity may produce similar scale variations. More broadly, our data suggest that other spatial covariates remain to be described, as some samples showed luminal regions with unique patterns of gene expression (fig. S9G) or low-DC1 expression signatures throughout the entire lumen (fig. S9, H to M).

To determine the operons that underlie the mucus-proximal adaptation of *B. theta* identified with DC1, we separated spatial patches into low-DC1 (mucus-associated) or high-DC1 (lumen-associated) compartments. We found 38 operons with statistically significant enrichment between these two compartments (Fig. 4L), with substantial overlap in the enriched operons when samples from the two fixation methods were analyzed separately (fig. S9, N to Q). Interestingly, there was a clear difference in the basic gene categories differentially expressed between the mucus- or lumen-associated compartments. 9 of the 15 lumen-associated operons were connected with central metabolism while none of the 23 mucus-associated operons had this functional annotation (Fig. 4L), suggesting that luminal *B. theta* may upregulate elements of central metabolism relative to *B. theta* closer to the mucus layer. By contrast, all 23 mucus-associated operons contain SusC or PUL-associated HTCS genes, as opposed to only 5 of the 15 lumen-associated operons, suggesting mucus-associated niches support harvesting of a greater polysaccharide diversity.

We next examined the known substrates for the PUL enriched in each compartment. Of the mucus-associated PUL with known substrates, 10 target host-mucus polysaccharides while only 1 targets dietary polysaccharides (Fig. 4L). By contrast, in the luminal compartment, 2 of the 3 PUL with known substrates target dietary polysaccharides (Fig. 4L). These observations are consistent with a greater availability of host-derived polysaccharides near the mucus, as expected, supporting our spatial analysis. Moreover, of the 23 PUL enriched in mucus-associated patches, 10 have unknown substrates (Fig. 4L). Given that we observe these operons enriched in mucus-associated patches, we predict that these PUL target host-derived polysaccharides or other carbohydrates enriched in this local environment. Our measurements complement a previous microdissection study (*63*) of the spatial variation of *B. theta* gene expression by providing a direct micron-scale measure of this variation and by extending the list of mucus-enriched PUL due, perhaps, to increased spatial resolution, improved sensitivity, or biological variability between the studies.

In total, these measurements reveal that bacteria fine tune gene expression to adapt to micron-scale niches in the gut. Excitingly, with bacterial-MERFISH it should now be possible to profile such micron-scale adaptation to a wide variety of complex environments.

## Discussion

Here we introduced bacterial-MERFISH, an image-based approach to single-cell transcriptomics that overcomes the massive mRNA density within bacterial cells by combining an optimized expansion microscopy toolbox with MERFISH. Bacterial-MERFISH offers complementary benefits to the growing suite of bacterial scRNA-seq methods (*5–14*). As a targeted method, it can sidestep abundant RNA (e.g., rRNA) challenges while providing an opportunity for high detection efficiency, which may prove essential for the many bacterial mRNAs that are very lowly expressed. In parallel, bacterial-MERFISH can be highly multiplexed, providing the ability to screen transcriptional changes with minimal prior knowledge of relevant targets. Bacterial-MERFISH can also image large numbers of cells, which may prove useful in the characterization of rare phenotypes. As an image-based technique, it naturally links gene expression to cell morphology or to intracellular molecular organization. Finally, with recent advances in all-optical readouts of pooled genetic screens (*65*, *66*), it may now be possible to combine transcriptome-wide profiling with genome-wide perturbations in bacteria.

However, we anticipate that one of the most substantial advantages of bacterial-MERFISH will be the ability to profile bacterial behaviors *in situ*. Whether it is interactions between specialized cellular states within single-species biofilms or between different species in mixed communities, bacterial-MERFISH provides a means of linking spatial proximity, cellular micro-environment, and global architecture to gene expression. Studying bacteria in their native environment also bypasses the need for culture; thus, bacterial-MERFISH may offer an avenue for the *in situ* characterization of the diverse range of unculturable bacteria. Finally, as MERFISH can now target both eukaryotic and prokaryotic mRNAs, bacterial-MERFISH may allow the simultaneous profiling of host and bacterial gene expression, which may deepen our understanding of host-microbe interactions such as commensal colonization or pathogenic infection. More broadly, the ability to directly profile single-cell bacterial transcriptomes in their native, complex environments may offer a new window into the substantial range of bacterial behaviors not well captured in a culture flask.

## Acknowledgements

We thank members of the Moffitt laboratory and S. Helaine for helpful discussion and a critical reading of the manuscript. We thank L. Bry, V. Yeliseyev, and M. Delany at the Massachusetts Host-Microbiome Center (MHMC) for providing monocolonized mice. Portions of this research were conducted on the O2 High Performance Compute Cluster, supported by the Research Computing Group, at Harvard Medical School. We thank the Dana-Farber/Harvard Cancer Center in Boston, MA, for the use of the Rodent Histopathology Core, which provided paraffin embedding and sectioning service. The Dana-Farber/Harvard Cancer Center is supported in part by an NCI Cancer Center Support Grant (P30CA06516). Access to this core and the MHMC were supported by the Harvard Digestive Disease Center (P30DK034854).

## Author contributions

Conceptualization: AS, YW, JRM Methodology: AS, YW, JRM Investigation: AS, YW, NTA, JRM Formal analysis: AS, YW, NTA, JRM Funding acquisition: JRM Supervision: JRM Writing – original draft: AS, YW, NTA, JRM Writing – review & editing: AS, YW, NTA, JRM

