## Supplemental Methods and Figures for "Highly Multiplexed Spatial Transcriptomics in Bacteria"

### Materials and Methods

*E. coli* Probe Design

To design a probe set that targets 101 operons in *E. coli*, we randomly selected operons to cover a range of expected abundance using published bulk RNA-sequencing data (*51*). MERFISH encoding probes are comprised of a target region which is complementary to the RNA of interest and flanking readout sequences. There is one readout sequence associated with each bit in the barcode assigned to targeted RNAs (*28*). To select the target regions for this library, we used a previously described pipeline (*67*) with the following constraints: 30-nt length; a melting temperature between 65 °C and 75 °C; a GC content between 40% and 74%; a gene specificity between 0.75 and 1; no predicted homology with rRNA; and no allowed overlap between target regions. For this initial library we chose to design target regions that would bind to a single gene within polycistronic operons. However, in a few instances, genes were too short, and we extended target region selection to the entire predicted operon. No final target had less than 54 target regions. The target region design pipeline is available at https://github.com/ZhuangLab/MERFISH_analysis.

To encode target identity, we assigned to each target a unique 16-bit binary barcode drawn from a set with a minimum Hamming distance of 4 and a constant Hamming weight of 4. Out of the 140 possible unique 16-bit barcodes satisfying our Hamming parameters, we used the 39 barcodes that were not assigned to any target mRNA as false-positive ('blank') controls. Each bit in the barcodes was then associated with a unique readout sequence (*67*, *68*). As our barcodes use a Hamming weight of 4, each mRNA target is, thus, defined by four unique readout sequences. To create the sequences of the final encoding probes for a given mRNA, we concatenated two of the four readout sequences associated with that RNA to each of its target regions. To create template molecules for the construction of these encoding probes, a T7 promoter and two unique PCR primers were concatenated. After the template probes were created for this library, we noticed two instances in which the genes we targeted are likely transcribed on the same operon (*infB* and *pnp*; and *bamA* and *dnaE*). As different barcodes assigned to the same transcript will likely be frequently corrupted, we chose to discard these four barcodes from all subsequent analysis, reducing this library effectively to a 97-operon library, which is how we refer to this library throughout.

For the design of the 1,057- and 1,930-operon libraries, we developed a more direct method for considering polycistronic operon structure. Specifically, we leveraged the operon structure annotations provided in Ecocyc v27.1 (*69*). Operons can have alternative transcription start and stop sites, leading to differences in the genes contained on polycistronic messages. As such variation raises the possibility that two different barcodes could be assigned to genes on the same message, we adopted a conservative approach to defining polycistronic messages. Specifically, we merged overlapping operon annotations to generate consensus sequences that did not share genes with any other consensus operon. We named the consensus operons based on the set of all the genes encompassed by the merged operon variants. Next, we used the same pipeline as described above to design target regions to these consensus operons with two notable exceptions. First, we excluded a handful of the most abundant mRNAs (e.g., some ribosomal protein mRNAs), which would have disproportionately affected target mRNA density. Second, we allowed some overlap in the potential target regions as some operons were too short to support 50 non-overlapping target regions. For the 1,057-operon library we allowed probe homology regions to overlap by 10 nt while for the 1,930-operon library we allowed target regions to overlap by up to 20 nt. In both cases, these libraries targeted all consensus operons which could accommodate at least 50 probes. However, we generated probes for more available target regions if an operon could support it. To encode the targets for the 1,057- and 1,930-operon libraries, we employed a 31-bit or a 40-bit barcoding scheme, respectively, with the same Hamming weight and distance constraints as described above. The encoding probes and their template molecules were also created as described above.

*B. theta* Probe Design

To design a 159-operon MERFISH library to profile the adaptation of *B. theta* to different niches in the mouse colon, we leveraged a published list (*70*) of SusC polysaccharide importers and hybrid-two-component systems (HTCS) associated with polysaccharide utilization loci (PUL). In addition, we included a random selection of operons associated with core elements of central metabolism, including glycolysis, gluconeogenesis, and fermentation. The same approach to target region design described above was used with a slight adjustment in the constraints on GC content (40% to 74%) and the overlap permitted for target regions (no overlap). We targeted individual genes within most operons but extended probe design to nearby genes in the same operon if the target gene could not accommodate enough probes. We designed at least 49 encoding probes for each gene with two readout sequences per probe. An 18-bit binary barcode set with the same Hamming properties as described above was used to define these 159 operons with 14 barcodes used as blank controls. Encoding probe templates were built for these target regions as described above.

Encoding Probe Synthesis

Encoding probe template libraries were purchased as complex Oligo Pools (Twist Biosciences) and amplified to create large quantities of single-stranded DNA encoding probes using protocols described previously (*28*, *71*). Briefly, template oligo pools were amplified with limited-cycle qPCR to create *in vitro* template molecules, which were used to create large quantities of single-stranded RNA via a high yield *in vitro* transcription with T7 RNA polymerase. Encoding probes were then created via reverse transcription of these RNA templates. The RNA templates and the reverse transcription primer (which contained a terminal ribonucleoside) were then removed via alkaline hydrolysis. Probes were purified by solid-phase reversible immobilization (SPRI beads; assembled as described previously (*71*)), and encoding probes were concentrated to a stock concentration of 1 mM with ethanol precipitation. Probes were stored as single-use aliquots at -20 °C.

smFISH Probes

All smFISH probes used for validation experiments in *E. coli* and *B. theta* were designed using the approach described for the 1,057-operon library for *E. coli* or the 159-operon library for *B. theta.* We designed at least 55 target regions per targeted operon and concatenated a single readout sequence to all target regions associated with a specific operon. smFISH probes were synthesized as oPools by Integrated DNA Technologies (IDT) and used for staining directly without amplification.

Ribosomal RNA Probes

We designed FISH probes to target and barcode the 16S ribosomal RNA (rRNA) of *E. coli* to provide dense staining of cells or, in the case of the diauxic shift measurements, to also distinguish cells harvested from multiple time points which were combined and measured in a single MERFISH measurement. These rRNA probes were based upon the common Eub388 probe sequence, which targets a region of the 16S rRNA largely conserved across all bacteria (*72*). We modified this sequence by extending the targeted sequence to 30 nt of homology to the *E. coli* 16S rRNA to allow hybridization in the same conditions as used for MERFISH encoding probes. This target sequence was concatenated to different readout sequences, and these probes were synthesized with a 5'-acrydite moiety to enable copolymerization in expansion gels (*35*). Additionally, these probes were synthesized with ribonucleotides to prevent their digestion during DNase treatments (RA83-RA48). We also designed a DNA version of the extended Eub388 probe without the acrydite for experiments that did not involve expansion microscopy (DA48). The rRNA probes were synthesized by IDT.

*E. coli* Culture, Fixation, and Permeabilization

For samples cultured in Luria-Bertani (LB) broth, a single colony of *E. coli* K-12 MG1655 (CGSG #7740, Yale Coli Genetic Stock Center) grown on an LB-agar plate was inoculated in ~5 mL LB Lennox medium (Fisher, AC612725000) to create an overnight culture. The next day this overnight culture was diluted 1:100 in 30 mL of fresh LB and grown to an optical density (OD600) of 0.32-0.34 for harvest. Unless otherwise specified, all bacterial liquid cultures were grown at 37 °C with shaking at 220 rpm.

For samples associated with the glucose-xylose diauxic shift, an overnight culture of *E. coli* was created by inoculating a single colony grown on an LB-agar plate into 5 mL MOPS minimal defined medium (MDM; Teknova, M2106) supplemented with 0.2% w/v glucose (Teknova, G0520). The following day this overnight culture was pelleted, the supernatant was removed, the pellet was resuspended in 2 mL of fresh MDM, and the OD600 was measured. This resuspended overnight culture was then diluted into 400 mL of MDM supplemented with 0.025% w/v glucose and 0.025% w/v xylose (Sigma, X3877) to a final OD600 of 0.004. This culture was grown, and small volumes (~2 mL) were drawn to monitor growth by their OD600. To harvest cells for MERFISH, a variable quantity of culture was collected for fixation with the volume adjusted (between 7 and 21 mL) such that samples harvested at different stages of the growth curve had comparable numbers of cells for the filtration and rRNA probe hybridization steps below.

For paraformaldehyde-fixed (PFA) samples, fresh 32% w/v PFA (Electron Microscopy Sciences, 15714) was added directly into the LB or MDM cultures to a final concentration of 4% w/v PFA, and the sample was fixed for 30 minutes at room temperature. The sample was then transferred into a 30 mL syringe and drawn through a 0.22 μm mixed-cellulose-ester syringe filter (Fisher, 09-720-004) to separate the cells from the fixation solution. The syringe was replaced with a new syringe, and reverse flow of 10 mL of 1×Phosphate Buffered Saline (PBS; Thermo, AM9625) across the filter was used to resuspend the cells. This solution was then pushed back through the filter to re-immobilize the cells, and this gentle wash was repeated for a total of three times. Upon the final elution from the filter, cells were resuspended in 3 mL nuclease-free water in a fresh syringe and transferred to a 50 mL tube containing 7 mL 100% ethanol (KOPTEC 200 Proof Ethanol; VWR 71001-866) for permeabilization. After 1 hour of incubation at room temperature, cells were pelleted, and the supernatant was removed. The pellet was then resuspended in 1 mL 1×PBS before proceeding to subsequent staining. For the diauxic shift experiments, samples collected at early time points were stored in 70% ethanol on ice until all other samples could be collected, fixed, and permeabilized as described above.

For methanol-fixed samples, the unfixed culture described above was pelleted, the supernatant was discarded, and the pellet resuspended in 100 μL water. The sample was fixed by adding 25 mL of 100% methanol (Sigma, MX0480) and incubating for 2 hours at room temperature. The sample was pelleted, the methanol removed, and the sample was resuspended in 1 mL of 1×PBS. This process was repeated for a total of three washes before the pellet was resuspended in a final 200 µL of 1×PBS before proceeding with subsequent staining.

Nuclease-free water for all protocols was generated by reverse osmosis (MilliporeSigma, Synergy UV) with a Biopack polisher (MilliporeSigma, CDUFBI001). Unless otherwise specified here and below, all bacterial pelleting steps were performed at room temperature via a 600×g spin for 10 minutes on a tabletop centrifuge.

Coverslip Preparation

40 mm-diameter coverslips (Bioptechs, 40-1313-03193) were used for holding samples throughout the expansion protocol or during imaging. When necessary, these coverslips were silanized as described previously (*73*). Briefly, coverslips were first cleaned in 1:1 37% HCl (Sigma, 258148) and methanol and then coated with an allyl-silane layer. The allyl moiety was selected as it incorporates into polyacrylamide gels, creating a covalent bond between the stabilization gel and the coverslip. Silanized coverslips were stored at room temperature in a desiccated environment prior to use.

In some cases, silanized or non-silanized coverslips were treated with poly-L-lysine (PLL) to enhance the adherence of samples to the coverslips. PLL-coated coverslips were prepared by washing coverslips in 70% ethanol, covering them with a solution of 0.1 mg/mL PLL (Santa Cruz Biotechnology, sc-286689), incubating at room temperature for 10 minutes, and then washing away the excess PLL twice with 1×PBS.

Mouse Preparation, Tissue Collection, and Sample Processing

All mice were used in accordance with animal care guidelines from the Harvard Medical School Standing Committee on Animals and the National Institutes of Health under a protocol approved by the Harvard Institutional Animal Care and Use Committee (IACUC).

We obtained mice monocolonized with *B. theta* from the Massachusetts Host-Microbiome Center. Germ-free C57BL/6 female mice were maintained in gnotobiotic isolators under a strict 12-hour light cycle and at a constant temperature (21 ± 1 °C) and humidity (55% - 65%). 10-week-old female mice were colonized with *B. theta* VPI 5482 by oral gavage ($4.8 \times 10^8$ colony-forming units [CFU]/mL) and maintained on a standard chow (LabDiet, 5021). Mice were euthanized with $CO_2$ asphyxiation followed by cervical dislocation.

The entire colon was rapidly dissected and fixed either with methacarn or periodate-lysine-paraformaldehyde (PLP). We pursued two different methods for fixation to confirm that the observed spatial patterns in *B. theta* gene expression were not fixation dependent, selecting these particular methods as they can better preserve the delicate mucus layer.

For methacarn fixation, the colon was immersed in 60% methanol, 30% chloroform (Sigma, C2432), and 10% acetic acid (Sigma, 695092) and incubated at 4 °C for 48 hours. The samples were then washed with 100% methanol for 35 minutes at 4 °C for a total of two times, then washed with 100% ethanol for 30 minutes at 4 °C for another two washes.

For PLP fixation, the colon was rapidly immersed in 2% w/v paraformaldehyde, 2 mg/mL sodium (meta)periodate (Sigma, S1878), and 0.075 M L-lysine (Sigma, L8662) in 1×PBS and incubated for 3 hours at 4 °C. These samples were then washed twice with 1×PBS, and then dehydrated with successive ethanol washes increasing in concentration from 50%, 70%, 95%, to 100%. Each wash was performed at 4 °C for 30 minutes, and the final 100% ethanol wash was performed twice. Samples prepared with either fixation method were stored at -20 °C pending further processing.

Samples were then embedded in paraffin and sectioned by the Rodent Histopathology Core at the Dana-Farber/Harvard Cancer Center. Briefly, the samples were incubated in xylene (VWR, 89370-088) for 1.5 hours at room temperature twice and infiltrated with paraffin (Leica, 3801340) for 4-5 hours at 60 °C, followed by embedding in a paraffin (Leica, 3801320) block. Paraffin blocks were then cross-sectioned to 5 μm slices, which were mounted on PLL-coated coverslips. The six measured samples were taken from multiple locations in multiple fecal pellets from two mice, each fixed with either methacarn or PLP. Coverslips were dried at room temperature and stored at 4 °C prior to expansion or smFISH staining.

<u>50X Expansion for Liquid Culture</u>

To hybridize the rRNA probes, permeabilized cells were washed twice in 1 mL RNA FISH wash buffer (30% v/v formamide [Thermo, AM9342] in 2×SSC [Thermo, AM9765]), and then resuspended in 50 μL RNA FISH hybridization buffer (30% v/v formamide, 10% w/v dextran sulfate [Millipore, S4030], and 1 mg/mL yeast tRNA [Thermo, 15401029] in 2×SSC) supplemented with the appropriate rRNA probe at a final concentration of 8 μM. Samples were incubated at 37 °C overnight to hybridize these probes. The next day, samples were pelleted, the supernatant removed, and the pellet was resuspended in 400 μL RNA FISH wash buffer. Samples were incubated 30 minutes at 47 °C to promote the melting of off-target probes, pelleted, and the supernatant was removed. This wash step was repeated a second time. For multiplexed samples, one additional wash with RNA FISH buffer was performed, and a similar number of cells from each time point, as judged by pellet size, were mixed. To remove the formamide carried over from the RNA FISH wash buffer, samples were pelleted, the supernatant removed, and the pellet was resuspended in 1×PBS. This wash was repeated a total of three times. Pelleting of samples in RNA FISH hybridization buffer was performed with a 600×g spin for 15 minutes given the increased viscosity of this buffer.

As the bacterial cell wall can resist expansion, the cell was digested prior to embedding in the expansion gel (*38*). Cells were pelleted and resuspended in cell wall digestion buffer (400 U/mL mutanolysin [Sigma, M9901] and 0.1 U/μL RNasin [Promega, N2615] in 1×PBS). The volume of the digestion buffer was adjusted to bring the concentration of the cells to an OD600 of ~1. 40 μL of cells were placed on a PLL-coated coverslip, which was placed in a Petri dish, and the solution was spread over a 1-2 $cm^2$ area with the side of a P1000 pipette tip while avoiding direct contact with the coverslip surface. The coverslip was placed in a humidified oven for 2 hours at 37 °C to perform the digestion, and then washed delicately with 1×PBS for a total of three times.

Our expansion gel recipe follows that introduced with the TREx method (*36*) with notable modifications in the handling of these gels for MERFISH. Briefly, single-use, 980-μL aliquots of expansion gel solution (1.1 M sodium acrylate, 2 M acrylamide [Bio-Rad, 1610140], and 60 ppm bis-acrylamide [Bio-Rad, 161-0142] in 1×PBS) were prepared and frozen at -20 °C. We found that the quality of commercially available sodium acrylate varied from lot to lot, and, as reported by other groups (*74*), we avoided sodium acrylate lots with a strong yellow or orange color or a cloudy appearance when dissolved in water as they produced inconsistent expansion. Due to these quality issues, we sourced sodium acrylate from multiple vendors (Sigma, 408220; or Santa Cruz Biotechnology, sc-236893). Like other groups (*36*), we also made sodium acrylate in house by neutralizing acrylic acid (Sigma, 147230) with sodium hydroxide (Sigma, 72068) to a pH range of 8-8.5 and adding water to adjust the final concentration of sodium acrylate to 38% w/v. We observed no obvious links between the quality of bacterial-MERFISH and the source of sodium acrylate; however, variations in the sodium acrylate source may have contributed to some variation in the degree of expansion (fig. S1, A and B).

To embed samples in expansion gels, a gel monomer solution aliquot was thawed and mixed with N'-tetramethylethylenediamine (TEMED; Sigma, T7024) and ammonium persulfate (APS; Sigma, 215589) to a final concentration of 0.10% v/v and 0.10% w/v, respectively, to initiate polymerization. Glass slides (Ted Pella, 260439) were coated with GelSlick (Lonza, 50640) according to the manufacturer's instructions. 200 µL of this initiated gel solution was then spotted onto the coated glass slides. Excess 1×PBS solution on the digested sample coverslips was removed by gently tapping the side of the coverslips on a Kimwipe (Kimtech, Kimwipes), and each coverslip was inverted onto an initiated gel solution droplet to embed cells within a thin film of gel between the coverslip and the glass slide. The samples were placed in a nitrogen chamber (Embrient, MIC-101), a damp Kimwipe was added to maintain humidity within the chamber, and the chamber was then sealed and purged with nitrogen. The chamber was then placed at 37 °C for 2 hours to allow the gels to polymerize in an oxygen-free environment. Samples were removed from the chamber, and the coverslips were gently detached from the glass slides. The gels were then cut with a rigid razor blade (WB Mason, ATSP591915) to an asymmetric shape to distinguish the orientation of the gels after expansion.

To further digest cellular contents that would restrict expansion, the gels—still attached to the coverslip—were transferred into 6 cm-diameter Petri dishes (VWR, 25384-092) and covered with 4 mL protease digestion buffer (8 U/mL proteinase K [New England Biolabs, P8107S] in 50 mM Tris [Thermo, AM9856], 1 mM ethylenediaminetetraacetic acid [EDTA; Thermo, AM9849], 0.5% v/v Triton X-100 [Sigma, T8787], and 0.8 M guanidine hydrochloride [Sigma, G7294]). The digestion was performed in a humidified oven at 37 °C overnight. The next day, the digestion buffer was removed, and the gels were quickly rinsed twice with nuclease-free water. The gels either detached from the coverslips during the digestion or, if still partially attached after digestion, were gently dislodged with a wash in nuclease-free water. The gels were then transferred into fresh 6 cm-diameter Petri dishes for expansion. To further expand the samples, residual buffer was removed, gels were covered in ~10 mL nuclease-free water, incubated for 20-60 minutes, the water was removed, and this wash process was repeated (typically 3-5 times) until the gel visibly stopped increasing in size.

Because expansion gels change size in different buffers, which would be incompatible with the buffer exchanges required for MERFISH, we embedded samples within an acrylamide stabilization gel that prevented this buffer-dependent size change (*35*). Water was removed from the Petri dishes, and gels were covered in ~5 mL of stabilization gel solution (4% v/v 19:1 acrylamide/bis-acrylamide [Bio-Rad, 1610144] in water) for 30 minutes at 4 °C. The gel solution was then replaced with ~5 mL of initiated stabilization gel solution (4% v/v 19:1 acrylamide/bis-acrylamide, 0.05% w/v APS, and 0.05% v/v TEMED in water) spiked with a 1:10,000 dilution of 0.1 µm-diameter carboxylate-modified orange-fluorescent beads (Thermo, F8800), which were added to serve as fiducials during MERFISH imaging. The samples were incubated for another 30 minutes at 4 °C to allow this activated gel solution to fully penetrate. Next, excess stabilization gel solution was removed, and the gels—infused with initiated stabilization gel solution—were placed onto a nylon mesh sheet (McMaster-Carr, 9318T46), which was itself placed on top of an acrylic cutting board covered with parafilm (VWR, 13-374-12). The nylon and parafilm provided a convenient support for the manipulation of fragile expanded gels. Each gel was then cut into ~4 mm × 4 mm squares with a rigid razor blade, and these squares were transferred, sample-side-up, onto a second parafilm-coated acrylic board. A silanized coverslip was then placed on top of each sample, and the sample and coverslip were placed in a humidified, nitrogen-purged chamber (as described above) and incubated at 37 °C for two hours to allow the gel to polymerize and crosslink to the coverslip. We noted that gel stabilization caused a 30-40% linear shrinkage of the expansion gels relative to the pre-stabilization expanded size likely due to the addition of charged polymerization initiators. For this reason, our 50X expansion protocol does not reach the same final degree of expansion reported for the TREx protocol (*36*). The stabilized samples were either immediately further processed with the following steps of the 50X expansion protocol, used as input for the following steps of the 1000X protocol (described below), or stored in 2×SSC at 4 °C for later use.

MERFISH requires the penetration of encoding probes into the gel as well as fluorescently labeled "readout" probes to label the readout sequences during MERFISH imaging. We found that thick gels substantially increase both the time required for this penetration and background fluorescence. Thus, we trimmed gels to a uniform, thin thickness to enhance the rate of probe penetration. Briefly, the silanized coverslips carrying the samples prepared above were placed on a flat RNase-free surface with the stabilized gel facing up and briefly washed with nuclease-free water for a total of three times. A 2 mil (~50 µm) thick slotted shim (McMaster-Carr, 9722K25) was placed around the gel to serve as a height spacer, and the gel was gently thinned with a thin flexible razor blade (Electron Microscopy Sciences, 72003-01) using the shim to set the height of the razor blade and the thickness of the gel.

During the development of bacterial-MERFISH, we found that very lowly expressed genes were measured more frequently with MERFISH than expected from bulk RNA-sequencing, producing an apparent false-positive rate higher than the false-positive rate we estimated from our blank control barcodes. In our experience, the blank controls are an excellent predictor of false-positive rates for MERFISH in eukaryotic systems. Thus, we reasoned that there must be an additional source of false positives in these bacterial samples. RNA FISH probe binding to DNA is not typically considered to be a possible source of false positives in unexpanded samples as the DNA duplex prevents the binding of probes; however, we surmised that during expansion some portions of the bacterial genome might be effectively melted due to topological constraints, allowing the binding of MERFISH encoding probes to the genome. This effect may not have been noticed in previous eukaryotic expansion-MERFISH (*32*, *35*) due, perhaps, to the modest degrees of expansion in those works, a differential propensity for topological-stretch-induced genomic melting in bacteria, or the greater fraction of low-abundance transcripts in bacteria relative to mammalian cells. To eliminate this source of false positives, we digested the genome in our samples. Briefly, excess water from the gel trimming step above was removed, and a hydrophobic barrier was drawn around gels with an Aqua-Hold PAP pen (Electron Microscopy Sciences, 71311). Gels were then briefly rinsed with water, the excess water was removed, and the gels were covered with 50 µL of DNA digestion buffer (2.4 U/µL murine RNase inhibitor [New England Biolabs, M0314L] and 0.1 U/µL DNase I in 1×DNase reaction buffer [Thermo, EN0521]). Samples were digested for 40 minutes at room temperature.

To hybridize MERFISH encoding probes, the DNA digestion buffer was aspirated, and the samples were washed with a high-salt hybridization-and-wash (HSHW) buffer (30% v/v formamide in 10×SSC) for 10 minutes at room temperature for a total of three times. To hybridize the probes, excess wash buffer was removed, and each gel was covered with 50 µL of HSHW buffer supplemented with 10 µM (97-operon library) or 100 µM (1,057- or 1,930-operon libraries) of the MERFISH encoding probe libraries constructed above. The samples were then hybridized in a humidified oven at 37 °C for 48-72 hours. After hybridization, the samples were washed in HSHW buffer at 47 °C for 30 minutes for a total of two times. Samples were then washed in 2×SSC for 2 minutes for a total of three washes before proceeding with imaging. We found that the increased salt concentration in these buffers, relative to the standard hybridization buffers used for MERFISH and smFISH (*67*, *73*), improved binding of the encoding probes. This improvement may be caused by the negatively charged expansion gel inhibiting the penetration of the negatively charged probes, and high salt may shield these charges.

1000X Expansion for Liquid Culture

To perform a second round of expansion to produce 1000X-expanded samples, 50X-expanded samples were taken after the formation of the stabilization gel and before thinning. These samples were first thinned as described above; however, as we found it easier to handle thicker gels during the second round of expansion, a thicker, 3 mil (~75 μm) shim was used. The thinned gels were then incubated in 300-400 μL of the same expansion-gel monomer solution described above initiated with 0.10% v/v TEMED and 0.10% w/v APS. The gels were gently shaken on an orbital shaker for 30 minutes at 4 °C or room temperature to allow the full penetration of the gel solution. Excess gel solution was removed, and this step was repeated a second time with fresh initiated gel solution. The samples on coverslips were then inverted onto a fresh 200 μL droplet of initiated gel solution spotted on a Gel-Slick-coated glass plate. These samples were transferred to a humidified chamber, which was then purged with nitrogen, and the gels were allowed to polymerize for 2 hours at 37 °C. The coverslips with the gel attached were then gently removed from the glass plate, and any newly formed gel outside of the original 50X expansion gel was removed with a razor blade. Gels were rinsed with nuclease-free water, expanded (without additional cell wall or protease digestion), and embedded in a second stabilization gel as described for the 50X expansion protocol. As fiducial beads were already incorporated into the first stabilization gel, such beads were not added to the second stabilization gel solution. Samples were either stored in 2×SSC at 4 °C before proceeding or were immediately thinned to 50 μm, DNase-treated, and stained with MERFISH encoding probes using the same protocol described above for the 50X samples.

Unlike the original iterative expansion microscopy protocols (*75*), this approach does not degrade the previous gels but rather expands them together with the second expansion gel. This form of iterative expansion was inspired by the ExpansionRevealing protocol (*37*), which showed that degradation of previous expansion and stabilization gels was not necessary for a second round of expansion.

smFISH Staining of Unexpanded *E. coli*

*E. coli* in LB or glucose-xylose MDM were fixed with PFA, permeabilized, and stained with FISH probes using the same protocols described above for ribosomal probe staining but with a final concentration of 2 μM of the smFISH probes and 1 μM of the rRNA probe (DA48). Cells were washed, pelleted, and resuspended in 1×PBS. Cells in 1×PBS were deposited on a PLL-coated coverslip prepared as described above, incubated for 1 hour at room temperature, and then washed with 1×PBS.

### 50X Expansion of Colon Samples

Colon slices prepared above were deparaffinized by heating the samples at 60 °C for 20 minutes followed by four washes of xylene (Sigma, 534056), each performed at room temperature for 2.5 minutes. The slices were then washed in 100% ethanol for 3 minutes at room temperature for a total of two times, then washed in 95% ethanol for 1 minute and then 70% ethanol for 1 minute. Sections were post-fixed on PLL-coated coverslips by incubating in 4% v/v PFA in 1×PBS at room temperature for 10 minutes, and the PFA was removed with two 1×PBS washes.

We made two notable changes to the 50X expansion protocol for colon slices. First, in early pilot work, we found that a PFA-treatment alone did not seem to efficiently anchor RNAs into the gel for paraffin-embedded samples; thus, we adopted a modified form of an RNA anchoring method developed previously (*76*, *77*). Second, we found it difficult to orient expanded samples without the clear landmark provided by the nuclei of the host gut; thus, we did not perform DNase treatment in these samples.

To create an RNA-reactive gel-crosslinking agent, we were heavily inspired by the MelphaX compound which combines melphalan (a bi-functional nucleic acid alkylating agent) with acryloyl-X (a gel-reactive acrylamide) to create a crosslinking agent that decorates nucleic acids with gel-reactive acrylamide groups (*77*). As Acryloyl-X is expensive, we reasoned that we could replace it with a much less expensive NHS-ester containing methacrylamide (MA-HNS; Sigma, 730300), creating a MelphaX-like crosslinker we term MelphaMA. To synthesize MelphaMA, an 8 mM melphalan (Cayman Chemicals, 148-82-3) stock in DMSO (Invitrogen, D12345) was mixed with a 100 mM stock of MA-NHS in DMSO in a 4:1 ratio, and the reaction was incubated at room temperature overnight with gentle shaking. As NHS reactions are efficient in DMSO and the MA-NHS was in excess of the melphalan, we assumed that the reaction proceeded to completion and treated the final solution as 6 mM MelphaMA. Single-use aliquots of MelphaMA were stored at -20 °C in a desiccated container.

To treat samples with MelphaMA, the slices were washed once in 20 mM MOPS pH 7.7 (Sigma, M9381) for 30 minutes at room temperature and then incubated with 50 µL of a 1:1 mix of this buffer with 6 mM of MelphaMA in a humidified oven at 37 °C overnight. The samples were then washed twice in 1×PBS for 5 minutes at room temperature. To digest the cell wall, the samples were covered with ~50 µL cell wall digestion solution (800 U/mL mutanolysin and 0.1 U/µL RNasin in 1×PBS buffer) and incubated for 2 hours at 37 °C. After digestion, samples were washed with 1×PBS for a total of three times.

To expand the samples, sections were washed with 200 µL tissue expansion gel monomer solution which contained the gel monomer solution described above supplemented with 0.01% w/v 4-Hydroxy-TEMPO (4-HT; Sigma, 176141), 0.20 % v/v TEMED, and 0.20% v/v APS for 20 minutes at 4 °C for a total of two times. 4-HT was added to this solution to further slow the polymerization and allow better penetration of the solution into the tissue slice (*34*). The gel was then polymerized, the sample digested with protease, the gel expanded and stabilized, and MERFISH and rRNA probes were hybridized using the protocol described above for liquid cultures.

### smFISH of Unexpanded Colon Samples

Colon samples were fixed, paraffinized, sliced, deparaffinized, and rehydrated as described above. smFISH staining was performed as described for *E. coli* samples with hybridization solutions that contained a total concentration of 1 µM per smFISH probe set and 1 µM of the

DA48 rRNA probe. As this probe was based on the eubacterial Eub388 probe, we found that it also labeled *B. theta* 16S rRNA despite a few nucleotides of mismatch with the *B. theta* 16S rRNA sequence. Unlike the cultured *E. coli* samples, we found higher degrees of background in smFISH imaging of colon slices; thus, after the hybridization and wash of smFISH probes, we used a previously described embedding and clearing approach to reduce background (*73*). Briefly, the samples were washed in a non-expanding gel solution (4% v/v 19:1 acrylamide/bis-acrylamide, 50 mM Tris-HCl, 300 mM NaCl [Thermo, AM9759], 0.03% w/v APS and 0.15% v/v TEMED) for 2 minutes, then inverted onto a GelSlick-coated glass plate with a 65 µL droplet of the same gel solution. This thin film of gel was polymerized for 2 hours at room temperature, and then the coverslip with the gel was gently removed from the glass plate, washed with 2×SSC, and digested in a clearing buffer (2% v/v sodium dodecyl sulfate [SDS; Thermo, AM9822], 8 U/mL proteinase K, and 0.25% v/v Triton X-100 in 2×SSC) at 37 °C overnight. The next day samples were washed with 2×SSC for 30 minutes at room temperature for each wash with a total of five washes and then stored in 2×SSC at 4 °C prior to imaging.

<u>MERFISH and smFISH Imaging</u>

All samples were imaged on an automated home-built microscope system as described previously (*71*). Briefly, this system is comprised of an epifluorescence microscope body (Nikon Ti-2) with a Celesta laser light engine (Lumencor, 90-10521), a 60× CFI PlanApo oil objective (Nikon), a 10× CFI PlanApo air objective (Nikon), and two CMOS cameras (Hamamatsu, ORCA Flash 4.0) with a twin-camera color splitter (Cairn, TwinCam). The sample was illuminated at 750 and 635 nm for MERFISH and smFISH signals; 750 nm, 635 nm, and 488 nm for rRNA probes; 555 nm for fiducial beads; and 405 nm for 4',6-diamidino-2-phenylindole (DAPI). Samples were mounted in a closed-flow chamber (Bioptechs, FCS2) with a 1 mm-thick flow gasket (Bioptechs, 1907-1422-1000). Buffers were programmatically delivered via a home-built flow system comprised of a peristaltic pump (Gilson, MINIPULS 3) and a daisy-chained set of valves (Hamilton, MVP).

The first set of readout probes complementary to the readout sequences associated with the first two bits were stained prior to loading the sample on the microscope. The storage buffer was aspirated from samples, and they were covered with 5 mL of readout hybridization buffer (10% v/v ethylene carbonate [Thermo, A15735-36] and 0.125% v/v Triton X-100 in 2× SSC) containing 3 nM of each of the fluorescently labeled readout probes. Samples were incubated for 30 minutes at room temperature in the dark with gentle orbital shaking, and then washed with readout hybridization buffer that did not contain readout probes for 20 minutes at room temperature for a total of two washes. For samples that were not DNase treated, the wash solutions were supplemented with 8 µg/mL DAPI (Thermo, D1306). Samples were then washed once with 2×SSC before imaging.

For experiments that require multiple rounds of readout staining, imaging, and removal, a flow cartridge was assembled containing readout hybridization buffers containing 6 nM (for MERFISH samples) or 3 nM (for smFISH samples) of the appropriate fluorescently labeled readout probes for each of the hybridization and imaging rounds. Once loaded onto the microscope, the sample was covered with an imaging buffer (4 µM Trolox-quinone (*78*), 0.5 mg/ml Trolox [Abcam, AB120747], 1:500 recombinant protocatechuate 3,4-dioxygenase [rPCO; OYC Americas, 46852004], and 5 mM protocatechuic acid [Sigma, 37580] in 2×SSC) designed to reduce photobleaching and enhance fluorophore brightness, and the desired fields-of-view (FOV) were imaged. 10 to 15 z-planes, separated by 1 µm, were imaged per FOV with the number of planes set by the degree of expansion. Once imaging was complete, the buffer was exchanged with a readout cleavage buffer (50 mM Tris(2-carboxyethyl)phosphine [GoldBio, TCEP25] in 2×SSC) and incubated, with gentle flow, for 15 minutes. This buffer was then exchanged with 2×SSC to remove residual cleavage buffer, and the next readout hybridization solution was added to the sample. The sample was incubated in the readout hybridization buffer for a total of 30 minutes, with gentle flow during the incubation, and then this buffer was exchanged with a wash buffer identical in composition to the readout hybridization buffer but lacking readout probes, in which the sample was incubated for 11.5 minutes. This buffer was then exchanged with the imaging buffer, and the sample was incubated in this buffer for 10 minutes prior to the start of the next imaging round to allow the rPCO to scavenge any oxygen introduced into buffers during flow. For *E. coli* samples, we performed 8 (97-operon library), 16 (1,057-operon library), or 20 (1,930-operon library) rounds of staining and imaging to identify all readout sequences. Where necessary, additional rounds of staining and imaging were performed to identify all utilized rRNA probes. For *B. theta* samples, we performed 9 rounds of staining and imaging for the 159-operon library. Readout probes, conjugated to Alexa488, Cy5, or Alexa750 via disulfide bonds, were synthetized by Bio-Synthesis.

<u>MERFISH Decoding</u>

To identify molecules from the raw MERFISH images collected above, we utilized a published analysis pipeline (https://github.com/ZhuangLab/MERFISH_analysis) (*67*, *68*). Briefly, this pipeline registered and aligned images taken of the same FOV from different imaging rounds with affine corrections built from the location of the fiducial beads to account for imperfect stage movements and affine corrections determined by multi-color samples to correct for small chromatic aberration in the imaging optics. Next, background noise was removed with a high-pass filter, and RNA signals tightened with Lucy-Richardson deconvolution. The intensity profile was then normalized across all imaging rounds, and the normalized intensity profile across all imaging rounds for each pixel was then compared via Euclidean distance to the profiles expected from all barcodes, and a pixel was assigned to a barcode if it was within a distance threshold set by a single bit flip. Adjacent pixels assigned to the same barcode were merged to create an identified RNA. Background signal was rejected by removing putative molecules with signal spread over too few pixels (the area) or with an average intensity too low (the brightness). In addition, molecules identified in the bottom-most z-plane were excluded from some datasets due to an increased false detection rate caused by nonspecific probe binding to the coverslip surface. For *E. coli* measurements, where we wanted to provide a conservative estimate of our detection efficiency, we identified likely instances in which the same RNA molecule was detected in different z-planes using DBSCAN to identify RNAs of the same type within 2 µm in the axial dimension and 0.2 µm in the lateral dimension and kept only the brightest molecule in each group.

<u>*E. coli* Bulk RNA-Sequencing</u>

To provide a validation of the mRNA abundance determined by bacterial-MERFISH, we collected bulk RNA-sequencing data. *E. coli* were cultured in LB or glucose-xylose MDM as described above. Samples were harvested at an OD600 of 0.33 (LB), 0.18 (glucose growth phase), or 0.47 (xylose growth phase). Total RNA was extracted using the RNAsnap protocol (*79*). Briefly, 4.5 mL of bacterial culture was mixed with 0.5 mL ice-cold stop solution (10% v/v phenol, pH 6.6 [Fisher, BP1750I-100] in ethanol), cells were pelleted with a 600×g spin for 8 minutes, and the supernatant was discarded. The pellet was then resuspended in 0.5 mL RNAsnap extraction buffer (9.5 mL 100% formamide, 360 µL 0.5 M EDTA, 25 µL 10% SDS, and 100 µL 100% 2-mercaptoethanol [Fisher, O3446I-100]), and the sample was incubated at 95 °C for 1 minute. Samples were stored at -20 °C as needed before proceeding. RNA was purified using the Zymo Direct-zol RNA kit (Zymo, R2051) by mixing 100 µL of the total RNA in the RNAsnap extraction buffer with 200 µL of the Zymo RNA binding buffer and then following the manufacturer's instructions.

Library preparation and sequencing for the MDM samples were performed by the Harvard Medical School Biopolymers Core using the Ribo-Erase kit for rRNA depletion, the KAPA HyperPrep kit for library preparation, and 150-bp paired-end sequencing on an Illumina NextSeq 500. Library preparation and sequencing for the LB sample were performed by Azenta using TURBO DNase for DNA depletion, the QIAGEN FastSelect rRNA HMR Kit for rRNA depletion, and the NEBNext Ultra II RNA Library Preparation Kit for Illumina according to the manufacturer's instructions. The library was 150-bp paired-end sequenced on an Illumina NovaSeq 6000.

Sequencing data were mapped to transcript abundance using salmon v1.8.0 (*80*). First, to facilitate comparison to the consensus operon structures used for the design of the MERFISH target regions, a salmon index was built on the sequences of these consensus operons. Salmon was then used, via the quant function, to measure the mapped count, effective length, and transcripts per million reads (TPM) for each operon. For consensus operons with disjoint probe design regions (e.g., when a noncontiguous subset of genes on the operon were targeted), we recomputed the TPM of the operon using the average mapped counts of contiguous regions weighted by their effective lengths.

MERFISH Cell Segmentation

To segment cells, an interactive segmentation toolkit, Ilastik 1.4.0 (*81*), was used to perform 3D pixel classification on the rRNA probe images. For computational efficiency, these images were downsized from 2048×2048 pixels to 512×512 pixels for 50X-expanded samples or to 256×256 pixels for 1000X-expanded samples. We used a human-in-the-loop Ilastik workflow to create a pixel classifier. For the LB measurements, this classifier classified pixels as in or out of a cell. For the diauxic shift measurements, pixels were classified as in a cell for a specific rRNA channel or out of a cell. The pixel classifiers for all data types were built using color, edge, and texture features: namely, we used the GaussianSmoothing, LaplacianOfGaussian, GaussianGradientMagnitude, DifferenceOfGaussians, StructureTensorEigenvalues, and HessianOfGaussianEigenvalues feature options provided by Ilastik with a series of 2D (1-40 pixels) and 3D sigma values (0.3-3.5 pixels). The range of the sigma values was tuned on a dataset-by-dataset basis based on the density of the rRNA stain, the image downsampling, and the degree of expansion for individual samples within the ranges provided. Ilastik was then used to predict class probabilities for the pixels of these downsampled images, which were then used to generate segmentation masks.

For 50X-expanded samples, preliminary masks were produced by thresholding on the pixel probabilities of the in-cell classification (>0.5), followed by morphological opening (with a radius of 1-2 pixels depending on the dataset) to separate contacting cells. These preliminary masks often had small internal regions of pixels that were below the in-class probability threshold. To fill these holes, a morphological closure operation (with a radius of 1-2 pixels depending on the dataset) was applied. This closure operation was applied to the mask of each cell separately to prevent the merger of adjacent cells. Contiguous regions within these masks were then identified and labeled as individual cells using scikit-image (*82*).

For 1000X-expanded samples the density of rRNA staining was not always sufficient to provide high-probability in-cell classification throughout the entire cell volume; thus, we adopted a seeded-watershed approach to define cell masks. To define the 'seeds', i.e. the locations in the image that most likely contain a cell, we used a dataset-specific, stringent in-cell probability threshold (0.4-0.7). To then define a foreground mask, we used a dataset-specific, less-stringent in-cell probability threshold (0.35-0.4). We then inverted the in-cell probabilities to create an image where high values correspond to regions of low cell probability. We next performed seeded-watershed on these inverted in-cell probability images in combination with the seeds and foreground masks using the watershed routine provided by scikit-image to create masks for individual cells.

Masks created for the 50X-expanded or 1000X-expanded samples were then filtered by volume (>60 pixels for 50X-expanded samples; >300-500 pixels for 1000X-expanded samples) and the integrated rRNA signal intensity (>1000 arbitrary units [AU] for 50X-expanded samples; >100 AU for 1000X-expanded samples) to remove spurious masks. This approach often produced mask boundaries that had a few-pixel roughness to the edge, as opposed to the smooth boundaries expected. Thus, a binary closure operation (1 pixel-radius for 50X-expanded samples or 5-pixel-radius for 1000X-expanded samples) was performed on the binary masks of each cell. To further smooth the edges of masks, the binarized mask of each cell was individually blurred with a Gaussian filter (sigma of 2), and a new binarized mask was generated by thresholding the values of this filtered image with a threshold of 0.2-0.3 for 50X-expanded samples, depending on the dataset, or 0.33-0.4 for 1000X-expanded samples.

For the diauxic-shift measurements with multiple rRNA channels, the Ilastik in-cell probabilities for each rRNA channel were processed separately as described above. This produced a set of segmentation masks for each channel, effectively associating each cell with a specific rRNA channel. However, in some instances the same pixels were assigned to multiple rRNA channel masks. To reconcile these conflicts and create consensus cell masks, these disagreements were settled by randomly selecting a priority for the rRNA channels and discarding overlapping masks from all but one channel based on this priority. To avoid preferentially favoring one channel over the others, this rRNA channel priority was randomized for each FOV.

RNAs were assigned to individual cells if they fell within the boundaries of the 3D masks as derived above. A small fraction of RNAs were found close to specific masks but just outside the boundary of these masks. Thus, we also assigned any mRNA within 1.2 μm (after expansion) of a mask to its nearest cell. When measuring the internal organization of the transcriptome, we noticed that outlier RNAs could have a disproportionate effect on the apparent coordinate system within each cell; thus, we adopted a more conservative distance threshold of 1.0 μm for these measurements.

<u>Estimation of Volumetric Expansion in *E. coli*</u>

To determine the degree of expansion in expanded *E. coli* cells, we started with 3D segmented masks generated as described above. For each mask, we measured the cell solidity and used it to eliminate segmentation artifacts or cells that were highly curved (which would challenge width measurements). Solidity was defined as the ratio of the volume of the segmentation mask to the volume of the convex hull that contains all the pixels within the mask. Segmentation artifacts or curved cells will have solidity values much less than 1, so, for this analysis, all cells with a solidity value less than 0.8 were removed. To define the width of cells, the mRNAs associated with each cell were projected onto a single 2D plane and the convex hull containing them was constructed. Given that the cells were oriented in various angles in space, cells were oriented by calculating a minimum bounding box around each convex hull. The dimensions of this box were then used to define a long axis which corresponded to the maximum length of the cell and a short axis which corresponded to the maximum width. As the poles can be narrower, we removed both poles by dividing the hull into four evenly spaced parts along the long axis and discarding the first and last quarters. The width of the cell was then estimated from the length of the intersection of lines perpendicular to the long axis with the boundaries of the convex hull. To estimate the width of each cell, multiple, perpendicular lines were measured at random positions along the central line, and the median of these values was used for the final width. To obtain an estimate of the linear expansion factor of each cell, the measured width was compared to a previous report of the width of *E. coli* MG1655 cell grown in similar conditions (1 µm) (*83*) (fig. S1A). To estimate the volumetric expansion factor, this linear expansion factor was cubed (fig. S1B). Finally, we calculated the mean expansion factor across 50X- and 1000X-expanded cells to obtain estimates of expansion by replicate.

Estimations of Bacterial-MERFISH Performance

To compare the average abundance per mRNA determined with bacterial-MERFISH to that determined via bulk RNA-sequencing, the Pearson correlation coefficient was computed between the $\log_{10}$ expression of each mRNA measured with both techniques. For datasets that were not segmented, MERFISH abundance was measured in average counts per FOV. For datasets that were segmented, MERFISH was measured in average counts per segmented cell.

As MERFISH is an image-based method, some cells may only be partially imaged. For example, cells that were aligned vertically or at an angle with respect to the coverslip would not be fully captured within the imaged z-stack. Similarly, cells that fell across the boundary of a FOV would also not be fully imaged. In parallel, errors in our segmentation masks can group two or more cells, effectively creating multiplet cells. For comparisons to bulk RNA-sequencing, all cells, including these partially imaged cells or potential multiplets, were retained in the analysis. However, to provide a measure of the detection efficiency, the number of mRNA copies, and the number of unique expressed operons per cell, we excluded potential multiplet cells and cells that were only partially imaged. For the 50X-expanded samples, cell multiplets were excluded by setting a minimum solidity threshold of 0.6, while partially imaged cells were identified as cells that were too small (less than 20 $\mu m^3$ mask volume), too close to a FOV boundary (within 120 pixels), or whose fraction of imaged cell volume within the top-most imaged z-plane was too large (more than 10% of mask pixels). For 1000X-expanded samples, we used a minimum mask solidity of 0.5, a minimum volume of 2000 $\mu m^3$, and a maximum of 7% pixels in the top z-plane. In both approaches, cells that were vertically aligned on the coverslip were identified and removed by requiring a minimum length to width ratio of 2, and occasional spurious masks were removed by eliminating all putative cells with less than 10 measured mRNAs. Finally, focus errors and large auto fluorescent debris can occasionally corrupt MERFISH measurements in some FOVs, leading to a dramatic reduction in the number of properly identified RNAs. Corrupted FOVs were identified from abnormally low numbers of detected mRNAs using the LocalOutlierFactor function of scikit-learn with the default parameters, and all cells within those

FOVs were excluded. Cells that passed all these cuts were deemed 'Whole' cells. Finally, the blank controls were not included in the number of detected mRNAs or operons per cell.

To estimate the detection efficiency of bacterial-MERFISH, published and calibrated bulk RNA-sequencing data (i.e., where abundance was measured in units of counts per cell) for *E. coli* grown in the same conditions was used to calibrate our own bulk RNA-sequencing data. First, the measured mRNA abundances in Bartholomäus et al. (*30*) (measured in reads per kilobase per million mapped reads) was converted to mRNA copy numbers by cell by scaling these reads by the ratio of the reported total copy number of mRNAs per cell (7,800) to the sum of the RNA abundances for all mRNAs. A conversion factor between the measured RNA abundances in our own bulk RNA-sequencing data (measured in TPM) to copy numbers per cell was calculated as the ratio of the total copy number per cell of all monocistronic operons in the Bartholomäus et al. measurements to the total TPM measured in our own data for the same operons. Bartholomäus et al. report abundances for genes instead of operons; thus, to avoid the challenge of properly estimating polycistronic mRNA abundances from those of the multiple genes they contain, the calculation of the conversion factor was restricted to monocistronic operons. The measured TPM of all the operons in our sequencing data were then multiplied by this conversion factor to obtain counts per cell. To determine the detection efficiency of MERFISH, we fit a line of slope 1 to the copy numbers per cell in each MERFISH measurement to the calibrated bulk RNA-sequencing abundances for the operons included in the measurement, using $\log_{10}$ expression to weight lowly and highly expressed genes more comparably. The detection efficiency was determined from the intercept of this line fit.

Analysis of *E. coli* Diauxic Shift

Three 1,057-operon MERFISH measurements were collected from two biological replicates of *E. coli* grown in glucose-xylose MDM, cells were fixed, 50X-expanded, rRNA-barcoded, measured with MERFISH, and segmented as described above. Counts per cell and cell metadata from all measurements were combined into a single anndata object (*84*) for downstream analysis.

To exclude segmentation artifacts and cells for which only a small fraction of the cell was measured, cells were filtered by volume (>25 $\mu m^3$), solidity (>0.6), MERFISH read counts (≥15 per cell), and number of unique operons detected per cell (≥7). Long cells tend to have lower solidity due to their increased curvature; thus, to better retain such cells, the minimum solidity requirement was relaxed to 0.5 for cells with a volume greater than 300 $\mu m^3$. The dimensions and orientation of each cell were identified from the principal components (PC) and eigenvalues of a principal component analysis (PCA) on the coordinates of the pixels within the masks. As cells are longer than they are wide, we found that the first PC aligned with the length of cells while the second and third PCs provided orthogonal measurement directions for the diameter of the cells. This analysis allowed us to further exclude cells with odd volumes given their lengths or with disagreements between two different estimates of the diameter, which are likely the result of segmentation artifacts. Specifically, cells were removed if their length (estimated from the first eigenvalue) to volume ratio was less than 0.35 or the ratio of the two diameter estimates (defined as the second eigenvalue over the third) was above 1.9.

The filtered anndata was then analyzed with the single-cell analysis toolkit scanpy (*84*) with default parameters unless otherwise specified. First, the number of operon counts per cell was normalized to a fixed target of 100 counts per cell, a pseudocount was added and these expression values were log normalized (with the natural base) and z-scored. Batch effects between the replicate MERFISH measurements were corrected with harmony (*85*) using the first

20 principal components as a basis. Differentially expressed genes were identified using the Wilcoxon rank-sum method on the $\log_2$-transformed normalized counts (with a pseudocount) without batch correction. A nearest neighbor graph with a neighborhood size of 20 was computed using the cosine distance metric on the first 20 harmonized PCs and was then used to compute a Uniform Manifold Approximation and Projection (UMAP) embedding (computed with a minimum distance of 0.2), a diffusion map, and to perform Leiden clustering with a resolution parameter of 1.3. Leiden clustering produced a total of 17 clusters, including three small clusters (<1% of all cells) which were excluded from subsequent analysis as they were enriched in blank counts and did not display strong differential expression of any operons. The reported marker genes for the Leiden clusters were selected from the top 15 genes per cluster with a minimum log fold change of $\log_2(2.5)$ and the highest marker significance scores reported by scanpy using the Wilcoxon rank-sum method.

To compute the pseudotime for cells transitioning from glucose log-phase growth through the diauxic shift, a cell in the late shift cluster N10 was selected as a starting point for the pseudotime trajectory, and the standard scanpy algorithm for diffusion pseudotime was applied. We chose to root the pseudotime analysis in cluster N10 as opposed to glucose log-phase growth due to the transcriptional heterogeneity seen in log-phase growth. For visualization purposes, pseudotime calculated in this fashion was then reversed to match the actual direction of the experiment from glucose into the diauxic shift. The second half of the pseudotime range was then monotonically rescaled to 0.9–1 with a piecewise-linear function to more equally weight the pseudotime ranges distributed across each diauxic condition. As we were interested in the response to glucose starvation, we did not include samples collected during the xylose-growth or stationary phase of the growth curve in this analysis. The reported operons for the pseudotime analysis were expressed in at least 3% of cells harvested in the remaining conditions and which contained at least one gene carrying the GO annotation of "carbohydrate metabolic process" or "carbohydrate transport" in Ecocyc v27.1.

<u>Analysis of the *E. coli* Transcriptome Organization</u>

The main analysis of the *E. coli* transcriptome organization was performed on cells taken from both replicates of the 50X-expanded, 1,057-operon MERFISH measurement of *E. coli* grown in LB; however, supporting analysis was also performed on 1000X-expanded samples. To further simplify this analysis, 'Whole' cells (as defined above) were further cut to retain only straight cells with sufficient RNA counts using a solidity threshold of 0.7 and a count per cell threshold of 40 for 50X-expanded samples, or 0.6 and 80 for 1000X-expanded samples.

To enable the comparison of intracellular RNA localization between different cells, a normalized cylindrical coordinate system was built for each cell, where each measured mRNA was localized by its position along the axial ($\boldsymbol{a}$) or radial ($\boldsymbol{r}$) axis. Formally, $\boldsymbol{a}$ and $\boldsymbol{r}$ were respectively defined as the distance from the RNA to the midcell along the long and short axes of the cell, normalized by the estimated length and diameter of the cell, respectively. We note that this definition ignores the distinction between 'old' and 'new' poles, which our measurements were not designed to identify, and assumes a radial symmetry to mRNA distributions.

To define this coordinate system for cells measured with 50X expansion, the orientation of each cell was determined using PCA on the 3D coordinates of the mRNAs associated with that cell, where PC1 indicated the long cell axis, and PCs 2 and 3 identified the radial plane. mRNAs were mapped to our intracellular coordinate system by projecting their 3D coordinates onto these three PCs and then normalizing by the dimensions of the cell. The cell dimensions were defined for each PC as the maximum projected distance between any two mRNAs in the cell. For each RNA, $\boldsymbol{a}$ was then given by the absolute value of the projection along PC1 and $\boldsymbol{r}$ by the L2 norm of the normalized projected distances on PCs 2 and 3.

Because of the lower final RNA density and relaxed solidity requirements in the 1000X-expanded samples, we found that the approach used for the 50X-expanded samples did not perform well. Thus, for the 1000X-expanded samples, we instead used the segmentation masks to define the intracellular coordinates. First, to correct for the differential resolution in X and Y versus Z, the segmentation masks were upsampled in Z to match the pixel size in X and Y. Each mask was then skeletonized with scikit-image to identify the central line of the cell. As skeleton endpoints are associated with the cell poles, cells with more than two end points in the skeleton were discarded as segmentation artifacts. This skeletonization process slightly eroded the cell central line from each of the two poles. To recover the original position of the poles, a second order polynomial was fit on the set of 10 pixels at each terminal end of the skeleton, and the intersection of these fit curves with the original cell mask was the used as an estimate of the pole location. The two polar intersection points as well as the pixel coordinates of the skeleton central line were then interpolated with a smooth spline, which was then resampled at equally spaced points to produce the final central line. To determine the values of $\boldsymbol{a}$ and $\boldsymbol{r}$ for each mRNA within the cell, $\boldsymbol{r}$ was defined as the Euclidean distance from that mRNA to the nearest point on the central line and $\boldsymbol{a}$ was defined as the integrated distance along the central line from this nearest point to the center of the cell. The length of the cell was measured as the full length of the center line, and the average radius of the cell was determined as the radius of a cylinder of cell length which would produce the volume of the cell segmentation mask.

To estimate the average distribution of individual operons, we used kernel density estimation (KDE). The volume of a cylindrical section depends non-linearly on the radius; thus, to create a volume-corrected 2D density projection, we first squared $\boldsymbol{r}$ to $\boldsymbol{r}^2$. To address boundary effects introduced by mapping mRNAs to the nonnegative quadrant of a symmetric cell, mRNAs were mirrored along the $\boldsymbol{a}$ and $\boldsymbol{r}$ axes. These mirrored mRNAs were then used to compute a KDE with scikit-learn and a bandwidth of 0.07. The KDEs were finally evaluated on 300 linearly spaced points along $\boldsymbol{a}$ and 100 linearly spaced points along $\boldsymbol{r}^2$, extending the evaluation range beyond the expected cell dimensions on each side by ~10% for visualization purposes. As the quality of the density estimates decreases when fewer observations are available, mRNAs with fewer than 500 measured molecules across all cells in both replicates were excluded.

To numerically compare different spatial patterns, the 2D densities described above were flattened into 1D vectors, these vectors were reduced in dimension to 150 components with PCA, and the pairwise cosine distances between different spatial patterns were calculated. These distances were then used to create a nearest neighbor graph using 15 nearest neighbors, which was then used, in turn, to create a UMAP embedding (with a minimum distance of 0.5) and to define Leiden clusters with a resolution parameter of 0.3. To determine the reproducibility of these spatial patterns, 2D KDEs for each mRNA were computed for each of the two replicates separately using the same approach. For this analysis, the minimum distance used to create the UMAP embedding was set to 0.25 and the Leiden clusters were defined with a resolution of 0.5.

To explore the correlation between the predicted location of the encoded proteins and the spatial patterns of mRNAs, PSORTdb 4.0 (*86*) was used to annotate *E. coli* operons with the predicted cellular location of the encoded protein. As polycistronic operons contain multiple genes which may have different protein locations, we defined the annotations of polycistronic operons as the union of the predicted locations of the individual genes they contain with one notable exception. In a co-translational insertion model of mRNA localization, co-translational insertion of inner-membrane-protein-encoding mRNAs should enrich such mRNAs at the membrane even if the mRNA contains genes that encode proteins that ultimately reside in other cellular compartments. Thus, in this model, a PSORTdb annotation of 'Cytoplasmic Membrane', which we redefine for clarity as 'Inner membrane', would supersede other annotations. Conversely, as proteins that are found within the periplasm or outer membrane are post-translationally transported, a co-translational-insertion model for membrane RNA localization would not predict membrane enrichment for mRNAs that encode such proteins. To test this model, we modified the annotations associated with each polycistronic operon: any operon that had at least one gene predicted to encode an inner-membrane protein had all other predicted locations for other genes discarded, whereas polycistronic operons that did not include a gene predicted to encode an inner membrane protein kept the full union of predicted locations in their annotations. While this model was applied to test the significance and enrichment of predicted encoded protein locations for the measured RNA localization clusters, all annotations are shown in fig. S6 for completeness. The significance and enrichment calculations were performed with the goatools package (*87*) using the Fisher exact test and a Benjamini-Hochberg false discovery rate (FDR) correction of 0.05 to define significant enrichments.

To compute the 1D axial distributions of the mRNAs, the radial component of the intracellular coordinates of the identified RNA molecules was discarded, the axial component was mirrored to enforce axial symmetry, and a 1D Gaussian KDE with a bandwidth of 0.2 was computed with scipy for each mRNA with more than 500 identified molecules across all cells. To explore how RNA localization may vary with cell cycle, we used cell length as a proxy for division stage and grouped cells by length. First, the pre-expansion length of the cells was estimated by dividing the length of their segmentation mask by their dataset-specific linear expansion factor computed above. Then, the cells were grouped by pre-expansion cell length in 0.6 μm increments and the axial KDEs were computed for each group separately. Because the number of cells in each group was sometimes too low to accurately estimate density for individual mRNAs, the mRNAs were binned by chromosome position in 10° increments (~130 kbp), and KDE was performed as described above on the chromosome bins instead of individual mRNAs.

To determine the spatial distribution of mRNAs measured with smFISH in unexpanded samples, where the spatial resolution was often insufficient to resolve individual RNA molecules, we developed an alternative approach to estimating average RNA localization. Specifically, 3D smFISH images were deconvolved, z-plane by z-plane, with a 2D Lucy-Richardson deconvolution with a Gaussian kernel of 7 pixels and a sigma of 2 pixels using the routines provided by scikit-image. In parallel, cells were segmented with Ilastik on the rRNA images as described above. Cells were then filtered based on the solidity (≥0.8), volume (≥50 voxels), planarity (the skeletonized mask must be contained within two z-planes), and length (a skeleton volume ≥ 7 pixels) of the segmentation masks. To eliminate cells that might span multiple FOVs, masks within 50 pixels of a FOV side were discarded. To align cells, PCA was performed on the 3D mask pixel coordinates to identify the orientation of the cell, and an affine transformation was applied to rotate both the mask and the deconvolved smFISH into a new coordinate system in which all cells are centered and have a uniform direction for the long axis of the cell. As cell cycle is correlated with cell length, cells were sorted into different length groups based on the length of the cell and resized with scipy such that all cells in the group had the same dimensions. The intensity profiles of all cells within a length group were then averaged to produce the final estimate of the spatial distribution of each mRNA measured with smFISH. The presented spatial distributions correspond to the average of cells post division, i.e., lengths between 3.3-3.8 µm, and were shown for a mid-cell z-plane.

Analysis of *Bacterial-MERFISH in the Mouse Colon*

Because expression levels for even abundant PULs can be low (fig. S8) and because the variation in *B. theta* density led to regional differences in the frequency of segmentation artifacts (when using similar Ilastik protocols as described above), we adopted a cell-free spatial analysis for the distribution of *B. theta* gene expression within the mouse colon. Moreover, as the density of mRNA expression varied substantially across regions within the colon and there was variability in expansion between samples, we adopted an approach that allowed for adaptive spatial binning. Specifically, we generated local patches of gene expression with fixed counts and dynamic size. A spatial patch was defined for each measured mRNA as the RNA itself plus its 49 nearest neighbors. To remove low mRNA density regions dominated by false positives, any patch that contained two mRNAs separated by more than 30 µm was discarded. By design, the patch construction process created partially redundant patches that could share many of the same mRNA molecules. Since making a non-redundant patch set is a form of the optimal packing problem, for which no computationally efficient exact solution exists, we applied a greedy algorithm to create a non-redundant set. In this approach, patches were selected at random and kept only if all mRNAs within that patch were unassigned to any other kept patch. The mRNAs within a kept patch were then marked as kept, and the process was repeated until all patches were either kept or discarded.

To eliminate patches disproportionately enriched in false positives, patches were further cut on the number of blank counts (greater than 3), the presence of unusually dim molecules (average mRNA brightness below the $2^{nd}$ percentile for all patches), or the presence of unusually bright molecules (average mRNA brightness above the $96^{th}$ percentile for all patches). The position of the host mucosa was determined via the intensity of the DAPI signal for all samples, and the distance between each spatial patch and the mucosal boundary was measured. The rare spatial patch that mapped to the host tissue was also removed. Based on visual inspection and the average count per cell determined from well segmented *B. theta,* individual spatial patches typically contained ~3-7 *B. theta* cells.

To identify reproducible spatial variation in the expression of *B. theta* in the colon, the spatial patches determined from four samples measured from a mouse prepared with the methacarn fixation and two samples measured from a mouse prepared with PLP fixation were combined in a single anndata structure, and a nearest neighbor graph of 15 nearest neighbors was created using a cosine distance metric. This graph was used to create a UMAP embedding with a minimum distance of 0.2 and to compute diffusion components (DCs) using the default parameters in scanpy.

To identify operons that were up or down regulated at low or high DC1 values, patches were categorized as low, high, or intermediate DC1 by dividing DC1 into 3 equally sized bins. The expression of each operon was averaged across all patches within each sample and DC bin, and the sample-averages were then used to calculate the $\log_2$-fold change in expression between the high- and low-DC1 bins. A t-test was performed with these sample-averages to determine the significance of this difference. The reported p-values were corrected for multiple hypothesis testing with the Benjamin-Hochberg FDR method as implemented in scipy. An FDR-corrected p-value threshold of 0.05 was used to define significance. The annotations of the target of each

PUL were sourced from dbCAN-PUL (*88*). In the rare instances in which a PUL listed multiple target polysaccharides with both host and dietary origins, we annotated the PUL as host-mucus-associated.

Utilized Public Software

MERFISH probe design and decoding were performed with public code (github.com/ZhuangLab/MERFISH_analysis) and were run in MATLAB r2021a. All other analysis was done in python 3.11.8 using scanpy (v1.9.8), anndata (v0.10.5.post1), harmonpy (v0.0.9), leidenalg (v0.10.2), umap-learn (v0.5.5), numpy (v1.26.4), scipy (v1.12.0), opencv-python (v4.9.0.80), scikit-image (v0.22.0), scikit-learn (v1.4.1.post1), tables (v3.9.2), tifffile (v2023.2.28), pandas (v2.2.1), and h5py (v3.10.0).

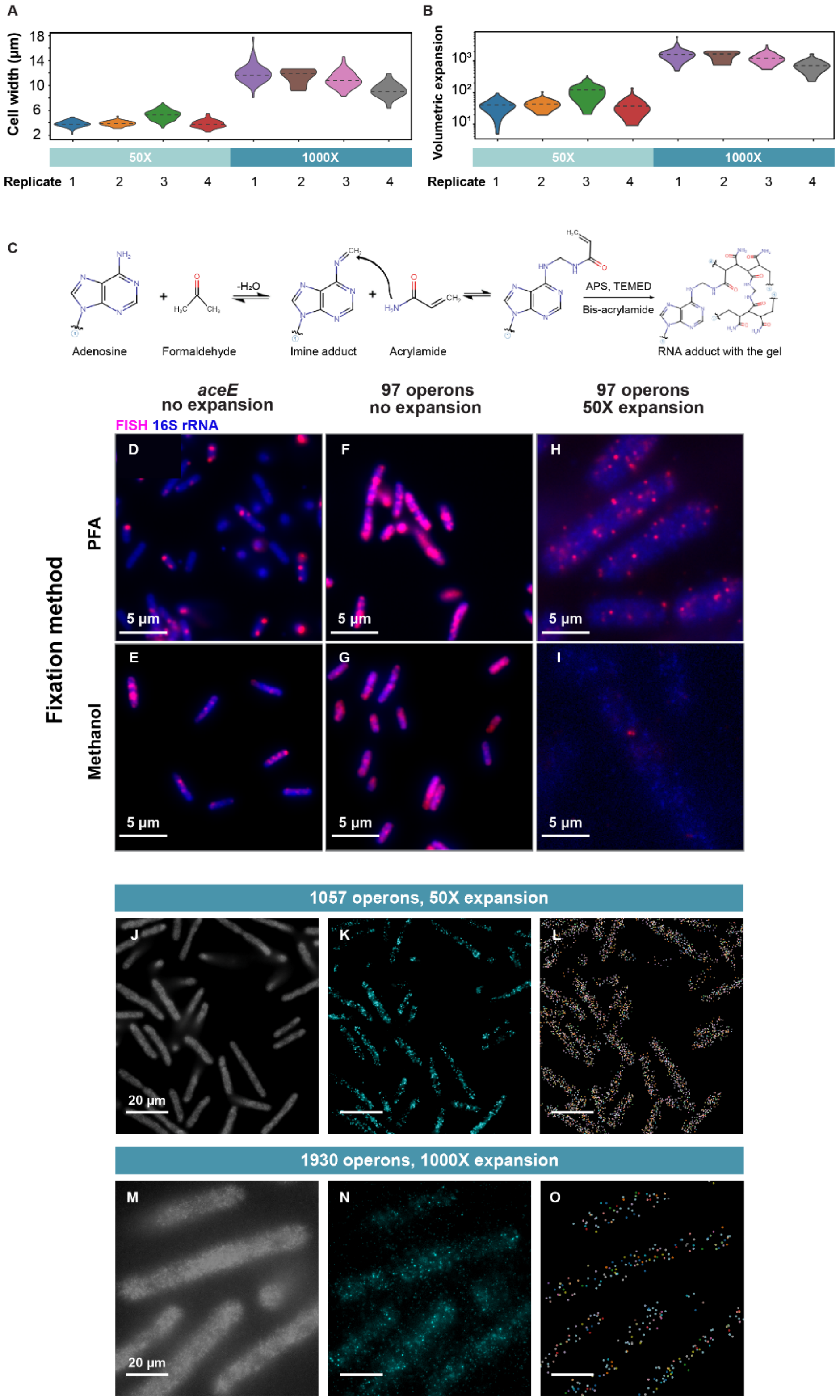


### Fig. S1.

**Properties of bacterial expansion and bacterial-MERFISH.** (**A**,**B**) Distribution of the measured width (A) or the estimated volumetric expansion (B) for multiple replicate measurements of log-phase *E. coli* grown in LB expanded with the 50X or 1000X protocol. Dashed lines represent the average. Distributions are kernel density estimates. The volumetric expansion is calculated from the cube of the ratio of the measured post-expansion width to the established 1.0-µm pre-expansion width (*83*). (**C**) A possible formaldehyde-based RNA-gel-anchoring mechanism that might explain the anchoring of RNAs to the expansion gel in the absence of specific RNA-gel anchoring chemistries introduced for eukaryotes (*76*, *77*). In this potential mechanism, formaldehyde adducts are created with RNA bases during fixation, and these long-lived adducts react with the amine groups in acrylamide moieties in the expansion gel, thus, anchoring RNAs to the gel. The full mechanism of RNA retention remains to be determined. (**D-I**) Images of log-phase *E. coli* grown in LB stained with a 16S rRNA probe (blue) and either smFISH probes against a single mRNA, *aceE* (D, E), or a 97-operon MERFISH library (F,G,H,I). *E. coli* were fixed either with paraformaldehyde [PFA] (D,F,H) or methanol (E,G,I). Scale bars: 5 µm. MERFISH images contain fluorescence only from mRNAs with a '1' in the first bit of their assigned barcode. FISH signals were comparable for PFA or methanol fixation prior to expansion but were dramatically reduced in methanol samples relative to PFA samples when expanded, supporting a PFA-dependent, RNA-anchoring mechanism. (**J,K**) Image of 50X-expanded log-phase *E. coli* grown in LB stained for rRNA (J) or the first bit of a MERFISH library targeting 1,057 operons (K). (**L**) The location and identity (color) of each RNA identified with MERFISH in the images in (J,K). (**M-O**) As in (J-L) but for a MERFISH measurement targeting 1,930 operons in 1000X-expanded samples. Scale bars: 20 µm.

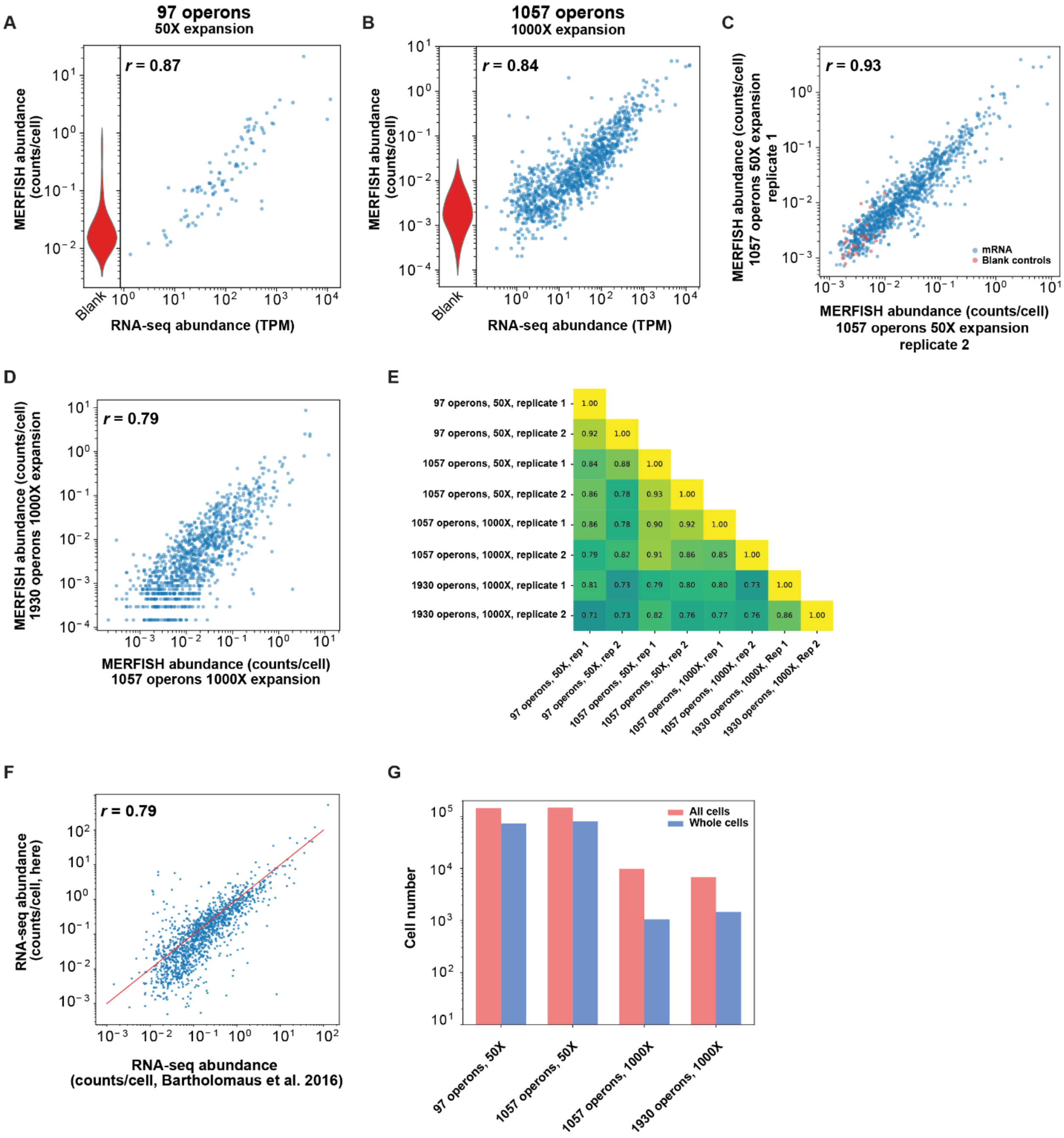


**Fig. S2.**

**Bacterial-MERFISH produces reproducible expression measurements in agreement with independent techniques.** (**A, B**) The average mRNA copy number per cell determined via MERFISH versus that determined by bulk RNA-sequencing (RNA-seq) of a matched culture for MERFISH measurements of 97 operons with 50X expansion (A) or 1,057 operons with 1000X expansion (B). Left: distribution of the false-positive ('blank') controls. (**C**) The average mRNA copy number per cell determined for one replicate of MERFISH of 1,057 operons with 50X expansion versus that of a second replicate. Blue markers represent mRNAs while red markers represent blank controls. (**D**) The average mRNA copy number per cell determined via MERFISH of 1,930 operons with 1000X expansion versus that determined via MERFISH against 1,057 operons with 1000X expansion. Only mRNAs shared between the two libraries are plotted. (**E**) All pairwise correlation coefficients between the logarithmic average mRNA copy number per cell determined for all combinations of MERFISH measurements against log-phase *E. coli* in LB for different numbers of targeted operons, different expansion protocols, and different biological replicates. (**F**) The average mRNA copy number per cell determined via bulk RNA-seq of log-phase *E. coli* grown in LB measured here versus that published by Bartholomaus et al. (*30*). The red line is equality. Bartholomaus et al. (*30*) calibrated their measured RNA abundances to produce counts per cell for each RNA, and we used those measurements to calibrate our own RNA-seq data (Materials and Methods). Where listed, *r* represents the Pearson correlation coefficient between the logarithmic expression values. (**G**) The total number of all (red) or whole (blue) cells measured across two replicates for each MERFISH measurement of *E. coli* in LB. Whole cells were conservatively defined as cells for which the entire volume could be imaged and which did not stretch across more than one field of view (FOV; Materials and Methods). All cells include whole cells as well as fractional cells that do not meet these criteria. All cells were used for abundance measures in correlation plots (Fig. 1, M to O and panels [A-E]). However, only whole cells were used for the calculation of detection efficiency, the number of unique operons per cell, and the total number of detected mRNAs per cell (Fig. 1, P to R). Because of the increased probability that a cell falls across an FOV boundary with the 1000X expansion, whole cells represented a smaller fraction of cells in those measurements as compared to 50X expansion.

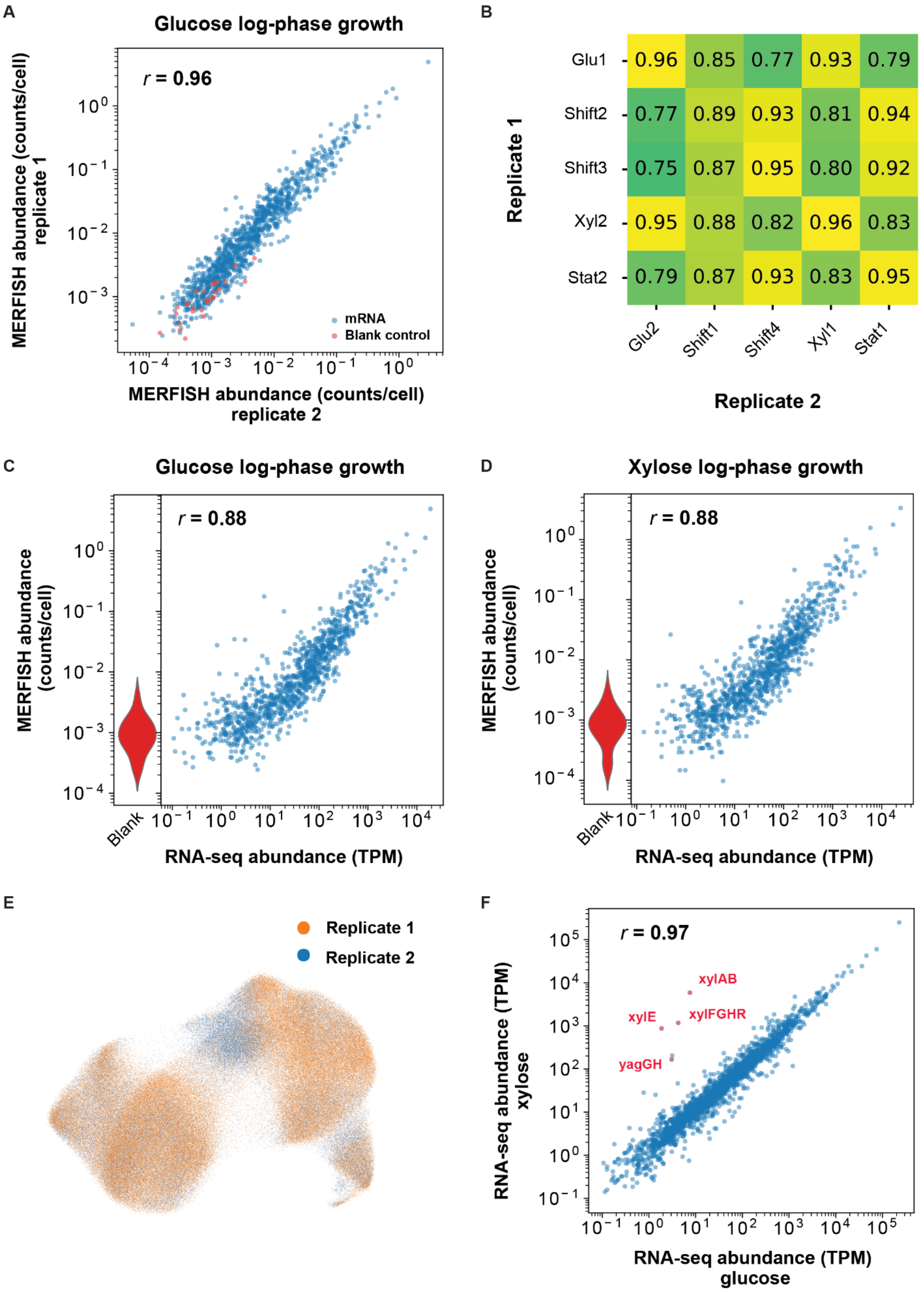
A
Glucose log-phase growth
r = 0.96
mRNA
Blank control
MERFISH abundance (counts/cell) replicate 1
MERFISH abundance (counts/cell) replicate 2
B
Glu1
Shift2
Shift3
Xyl2
Stat2
0.96 0.85 0.77 0.93 0.79
0.77 0.89 0.93 0.81 0.94
0.75 0.87 0.95 0.80 0.92
0.95 0.88 0.82 0.96 0.83
0.79 0.87 0.93 0.83 0.95
Glu2
Shift1
Shift4
Xyl1
Stat1
Replicate 1
Replicate 2
C
Glucose log-phase growth
r = 0.88
MERFISH abundance (counts/cell)
Blank
RNA-seq abundance (TPM)
D
Xylose log-phase growth
r = 0.88
MERFISH abundance (counts/cell)
Blank
RNA-seq abundance (TPM)
E
Replicate 1
Replicate 2
F
r = 0.97
xylAB
xylE
xylFGHR
yagGH
RNA-seq abundance (TPM) xylose
RNA-seq abundance (TPM) glucose

**Fig. S3.**

**Bacterial-MERFISH measurements in a diauxic shift agree with independent techniques and are reproducible.** (**A**) The average mRNA copy number per cell determined via MERFISH during growth in glucose for one replicate versus that measured in a second biological replicate. Blue markers represent mRNAs, and red markers represent false-positive blank controls. (**B**) Pearson correlation coefficients for the logarithmic expression of bacterial MERFISH measurements from different replicates and different time points in the glucose-xylose diauxic shift. (**C,D**) The average mRNA copy per cell determined via MERFISH versus the abundance determined via bulk RNA-seq during log-phase growth in glucose (C) or xylose (D). Left: distribution of blank controls. (**E**) UMAP of cells measured via MERFISH during the diauxic experiment as in Fig. 2 colored by the biological replicate in which each cell was measured. (**F**) mRNA abundance determined via bulk RNA-seq during log-phase growth in xylose versus that determined during log-phase growth in glucose. Individual operons associated with xylose utilization are marked. Where listed, $r$ represents the Pearson correlation coefficient between the logarithmic expression values. TPM: transcripts per million reads.

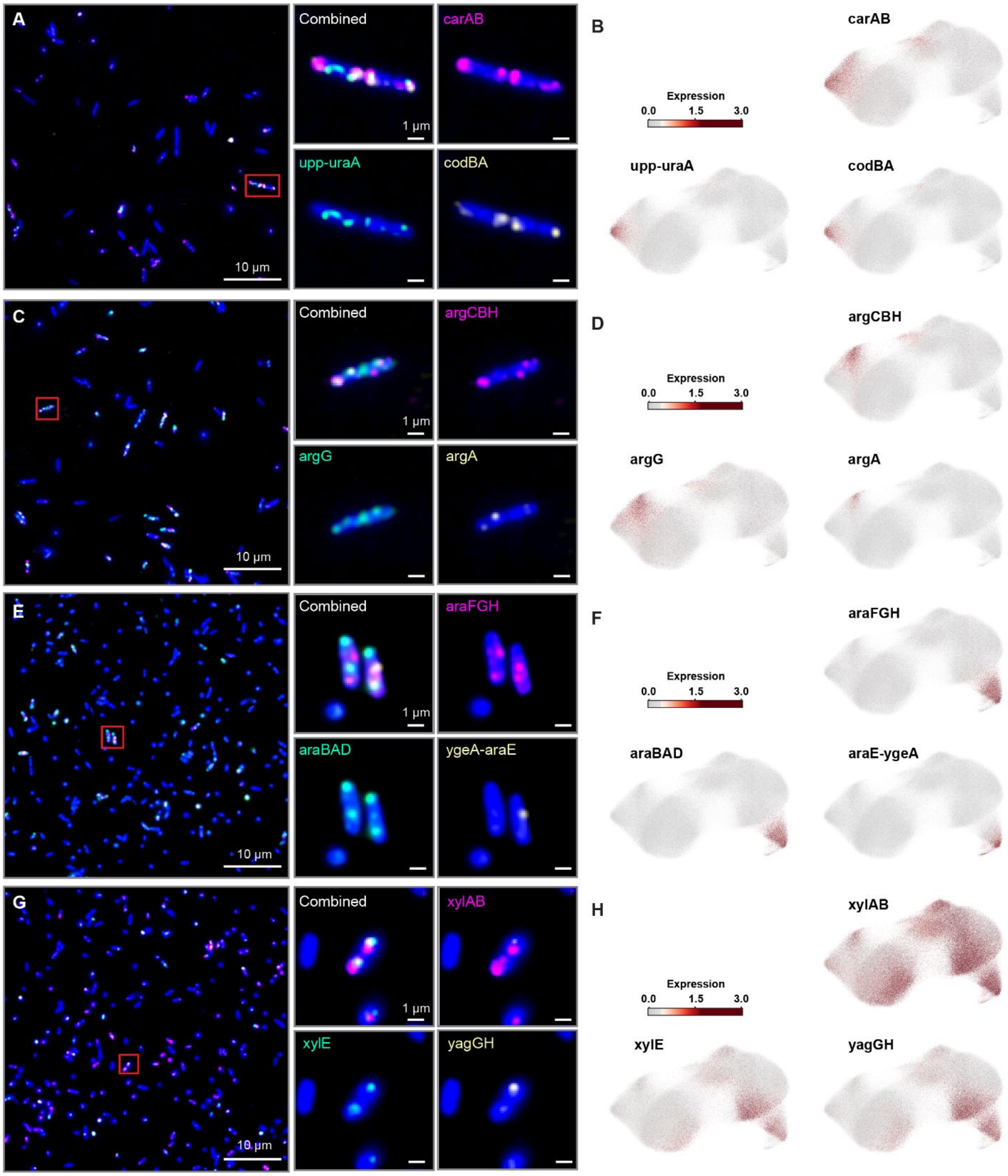


**Fig. S4.**

**Unexpanded smFISH supports clusters identified with bacterial-MERFISH.** (**A**) Images of unexpanded *E. coli* grown in a minimal defined medium with a mixture of glucose and xylose, fixed and stained for genes co-expressed in specific sub-populations identified with bacterial-MERFISH in the glucose-xylose diauxic shift. 16S rRNA stain is marked in blue, and smFISH for three different genes are colored in magenta, green, and yellow for both a large field-of-view (left) or the zoom-in boxed in red (right). Scale bars: 10 µm (left) or 1 µm (right). (**B**) UMAP colored by the natural log of the normalized expression of the listed genes. (**C,E,G**) As in (A). (**D,F,H**) as in (B). Co-expression of markers *carAB*, *upp-uraA*, and *codBA* define cluster G6; co-expression of markers *argCBH*, *argG*, and *argA* define cluster G7; co-expression of markers *araFGH*, *araBAD*, and *araE-ygeA* define cluster N10; and co-expression of markers for xylose utilization (*xylAB*, *xylE*, and *yagGH*) are associated with cluster N2. Cells in (A,C) were harvested during the glucose log-phase while cells in (E,G) were harvested during mid-to-late diauxic shift.

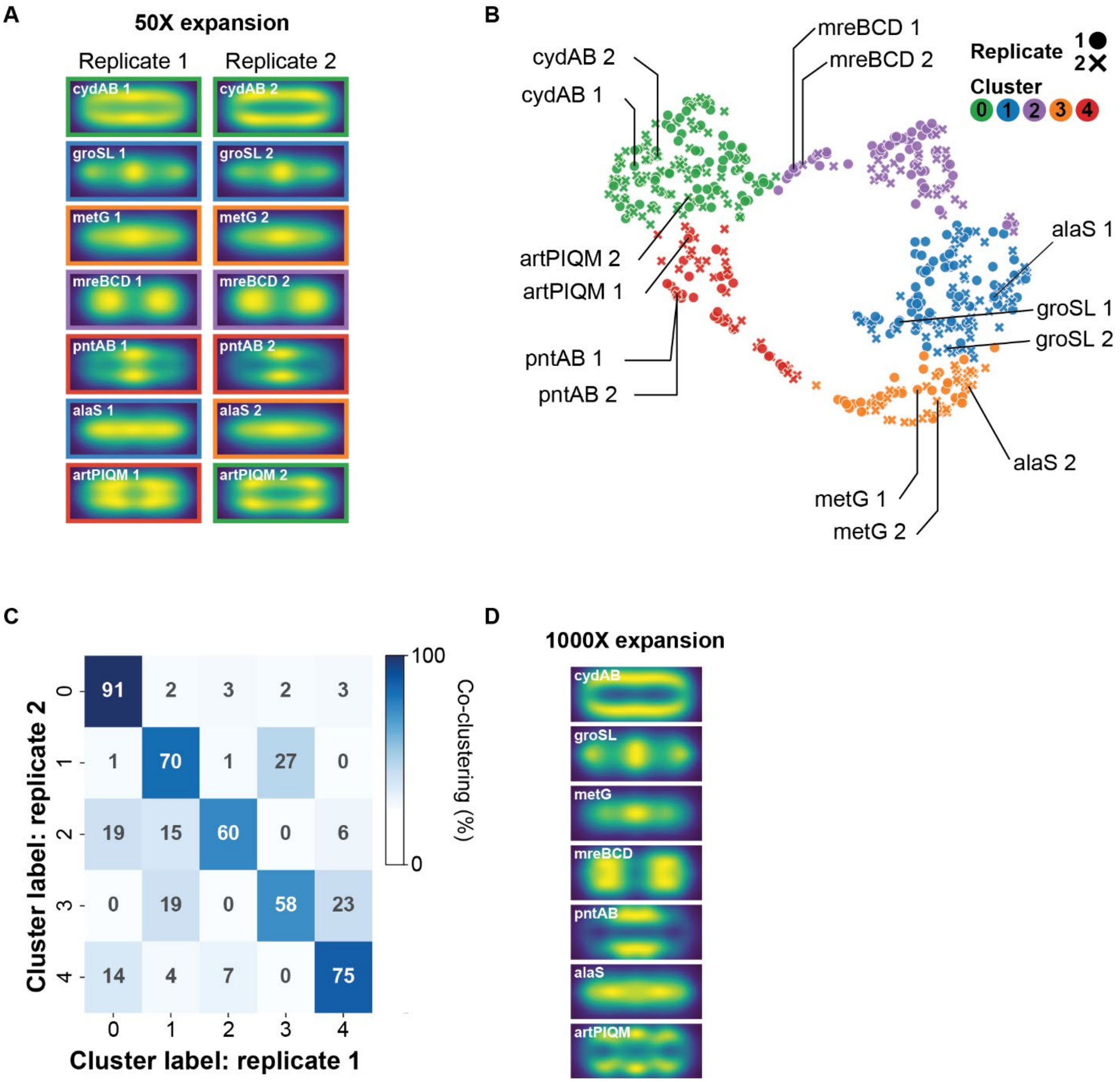


**Fig. S5.**

**Intracellular mRNA distributions are reproducible across replicates and techniques.** (**A**) Normalized mRNA density for MERFISH measurements of 1,057 operons determined in two replicates of 50X expansion. (**B**) UMAP of the spatial distribution of mRNA patterns with the replicates in (A) analyzed separately. Marker shape indicates replicate and color indicates Leiden cluster. As replicates were treated separately here, these clusters are distinct from those in Fig. 2. Representative operons are highlighted for each of the two replicates. (**C**) Fraction of mRNAs assigned a given cluster label (from panel B) from replicate 2 and assigned a given cluster label in replicate 1. The strong diagonal supports the reproducibility of our measurements and classification of spatial patterns. (**D**) Normalized mRNA density for MERFISH measurements of 1,057 operons with 1000X expansion for the same operons in (A).

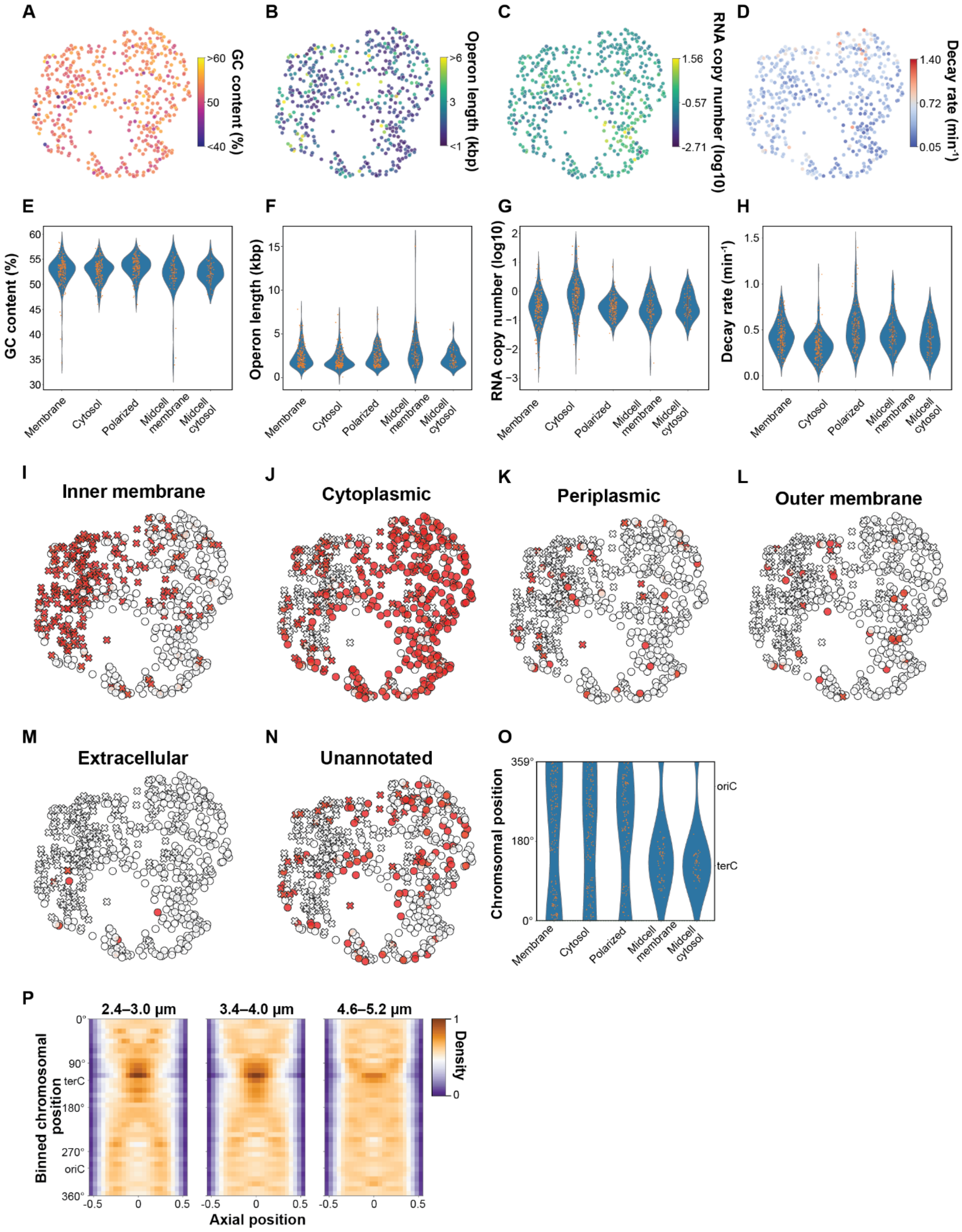


**Fig. S6.**

**mRNA features that covary with spatial patterns.** (**A-D**) UMAP of the spatial distribution of the *E. coli* transcriptome colored by GC content (A), operon length (B), abundance (C), and published mRNA decay rate (*51*) (D). (**E-H**) Distribution of GC content (E), operon length (F), abundance (G), and mRNA decay rate (H) for all mRNAs within each of the listed clusters. Markers represent individual mRNAs and the distributions were created with kernel density estimation. (**I-N**) UMAP of the spatial distribution of the *E. coli* transcriptome colored by the annotated location of at least one of the genes on each operon as determined by PSORTdb (*86*). Markers indicate whether any of the genes on that operon contain a gene encoding an inner-membrane protein (crosses) or not (circles). (**O**) Distribution of the location of the encoding chromosomal loci for mRNAs in each of the spatial clusters. The kernel density estimations were calculated recognizing the circular nature of the genome. The location of the replication origin (*oriC*) and terminus (*terC*) are listed. (**P**) The average axial mRNA distribution averaged across all mRNAs transcribed from chromosomal regions within defined position bins. Individual heatmaps represent the average for all cells that fall within the specified pre-expansion length range. This length serves as a proxy for cell-cycle stage.

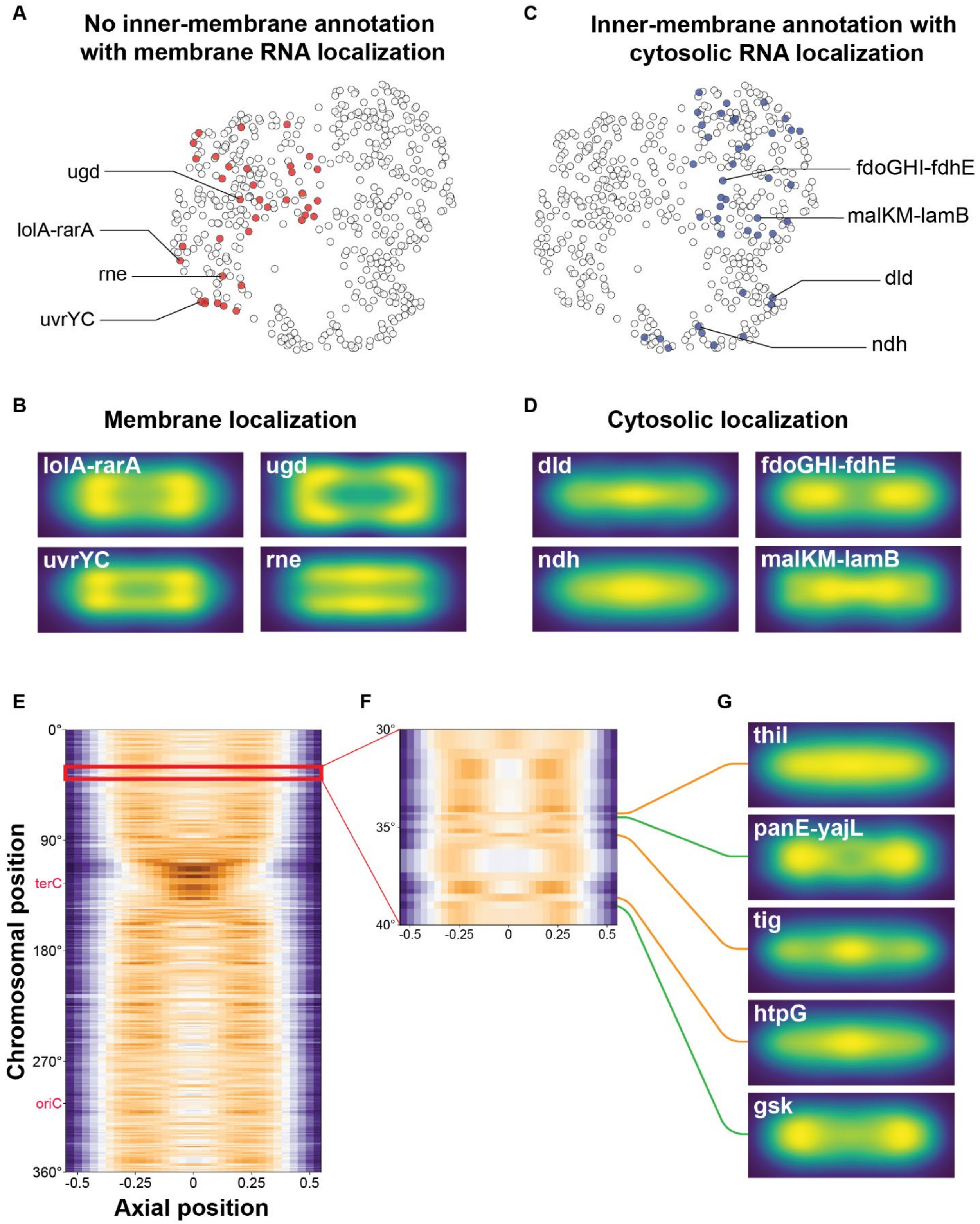


**Fig. S7.**

**Many mRNAs have spatial distributions that are exceptions to global patterns of mRNA localization.** (**A**) UMAP of the spatial distribution of the *E. coli* transcriptome highlighting (red) mRNAs that do not encode an inner-membrane protein and, thus, would not be predicted to be enriched at the membrane yet which were assigned to localization clusters with membrane-associated patterns. (**B**) Normalized mRNA density for MERFISH measurements of 1,057 operons with 50X expansion highlighting some of the exceptions listed in (A). (**C,D**) As in (A,B) but for mRNAs that do encode an inner-membrane protein and would be predicted to be found at the membrane yet were assigned to localization clusters with cytoplasmic or polar patterns. (**E**) The average axial distribution of mRNAs plotted by the chromosomal location of the encoding loci (reproduced from Fig. 3). (**F**) Zoom in on the boxed region in (E). (**G**) Spatial distribution of example mRNAs that do (green line) or do not (orange line) follow the predicted spatial patterns based on the location of their genomic loci.

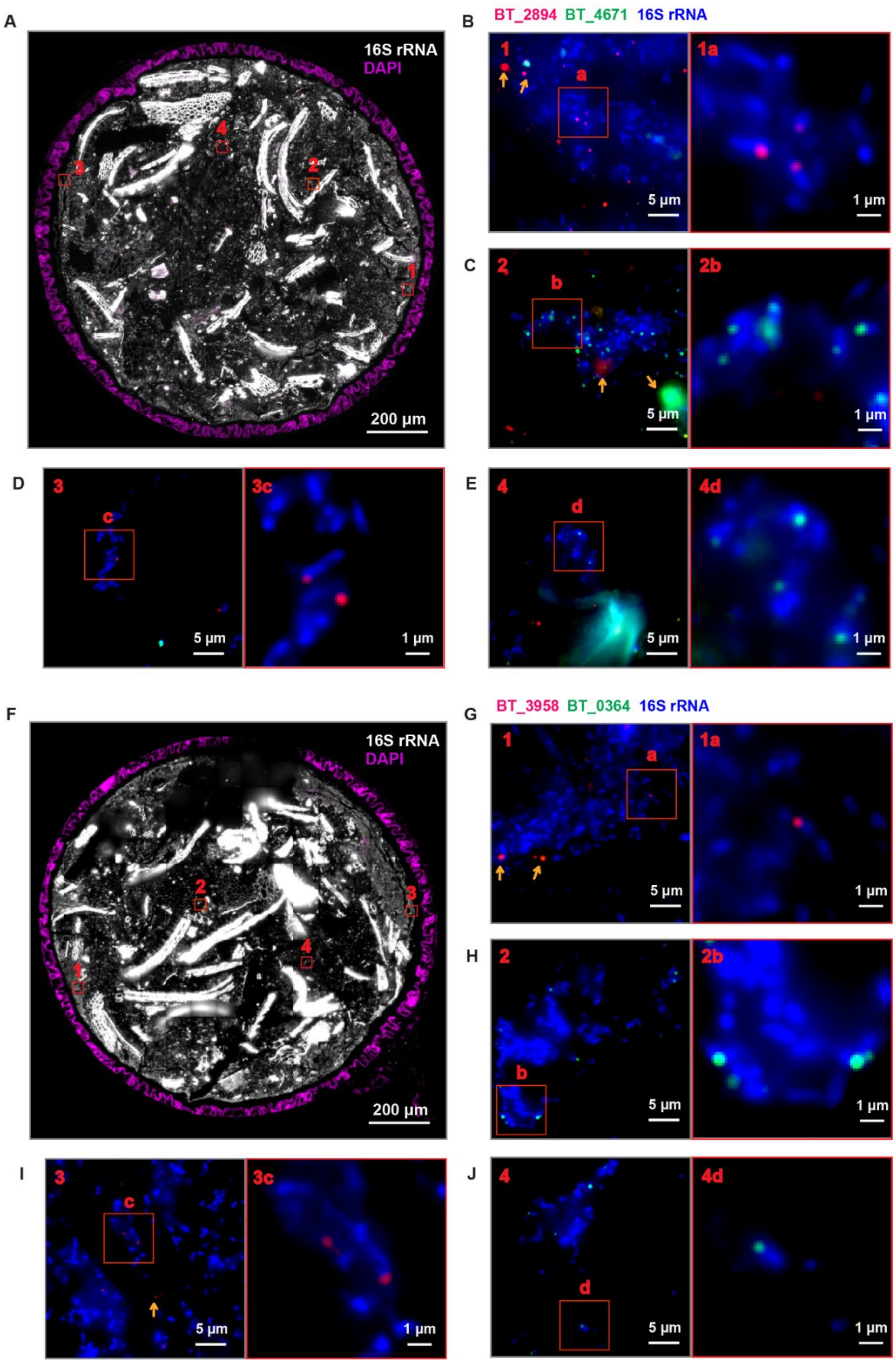


**Fig. S8.**

**smFISH measures spatial variations in the expression of several *B. theta* operons that are consistent with MERFISH.** (**A**) Image of an unexpanded colon cross section of a *B. theta* monocolonized mouse stained with DAPI (purple) and a 16S rRNA probe (grey). Scale bar: 200 µm. (**B-E**) Left: zoom-in on the slice in (A) stained with smFISH against two *B. theta* operons, BT_2894 (red), BT_4671 (green), and the 16S rRNA probe (blue). The number in the upper left indicates the red-boxed region in (A) represented in each image. Right: zoom-in on the lettered red box in the left panel. Arrows indicate regions of autofluorescence that should not be interpreted as mRNA signals as they do not overlap with rRNA signal or are too large to be single molecule signals. Scale bars: 5 µm (left) or 1 µm (right). (**F-J**) As in (A-E) but for a colonic section stained for BT_3958 (red) or BT_0364 (green). As seen with MERFISH in Fig. 4G, BT_4671 and BT_0364 are found expressed in the lumen while BT_2894 and BT_3958 are found preferentially expressed closer to the mucus layer. We also note that these genes, which are some of the more abundant of the *B. theta* operons probed, are expressed at very low levels such that only a subset of cells have detectable expression.

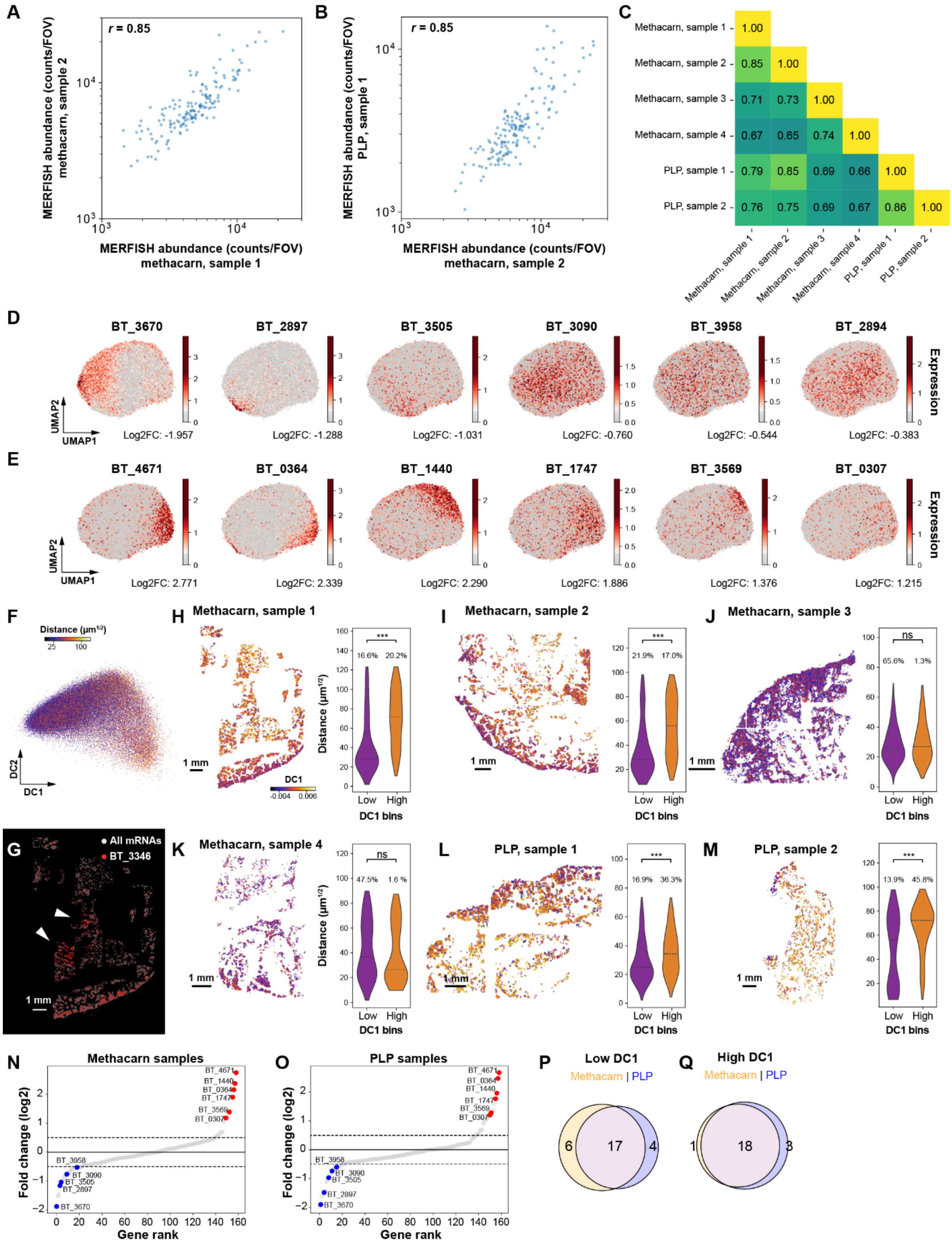


**Fig. S9.**

***B. theta*** **spatial expression patterns are reproducible.** (**A**) Average mRNA counts per field-of-view (FOV) determined for one colonic cross section fixed with methacarn versus that of another sample fixed in methacarn. (**B**) As in (A) but for a sample fixed with periodate-lysine-paraformaldehyde (PLP) versus methacarn. *r* represents the Pearson correlation coefficient between the logarithmic expression values. (**C**) Pairwise Pearson correlation coefficients for the logarithmic expression measured for all colonic cross sections of monocolonized mice. (**D,E**) UMAP of the local patches of *B. theta* colored by the expression of operons enriched at low DC1 (D) or at high DC1 (E). (**F**) Scatter plot of the first two diffusion components (DC1, DC2) for local patches of *B. theta* colored by the measured distance to the host tissue. Of the first two DCs, DC1 correlates most strongly with distance. (**G**) Spatial distribution of all RNAs (gray) or BT_3346 (red) for an example colonic slice. Arrowheads mark regions of expression of BT_3346 not seen in other regions of the lumen. (**H-M**) Spatial distribution of all measured *B. theta* patches colored by DC1 (left) or probability distributions for the distance of each patch to the host tissue for low and high DC1 bins (right) for each of the measured samples. Lines in the probability distributions represent median values. The listed percentages represent the fraction of patches within each DC1 bin. *** indicates a p-value of less than 0.001 as determined by a two-sided Wilcoxon test between the two distributions and 'ns' represents $p>0.05$. Scale bars: 1 mm. Collectively the results in (G-M) suggest that there is greater spatial heterogeneity in *B. theta* gene expression than described here. (**N,O**) The fold change ($\log_2$) for the expression of operons in high versus low DC1 patches sorted from smallest to largest fold change when just the methacarn-fixed (N) or PLP-fixed (O) samples are analyzed. Highlighted operons were identified when all samples were analyzed jointly. Dashed lines represent a $\log_2$ enrichment or de-enrichment of 0.5. (**P,Q**) Venn diagram showing the overlap in the low-DC1-enriched (P) or high-DC1-enriched (Q) operons between samples fixed with methacarn or PLP. Enriched genes were defined as those that exceeded the 0.5 thresholds in (N,O). The observed overlap supports the fixation-method-independence of these spatial enrichments.
